# Modeling and dissociation of intrinsic and input-driven neural population dynamics underlying behavior

**DOI:** 10.1101/2023.03.14.532554

**Authors:** Parsa Vahidi, Omid G. Sani, Maryam M. Shanechi

## Abstract

Neural dynamics can reflect intrinsic dynamics or dynamic inputs, such as sensory inputs or inputs from other regions. To avoid misinterpreting temporally-structured inputs as intrinsic dynamics, dynamical models of neural activity should account for measured inputs. However, incorporating measured inputs remains elusive in joint dynamical modeling of neural-behavioral data, which is important for studying neural computations of a specific behavior. We first show how training dynamical models of neural activity while considering behavior but not input, or input but not behavior may lead to misinterpretations. We then develop a novel analytical learning method that simultaneously accounts for neural activity, behavior, and measured inputs. The method provides the new capability to prioritize the learning of intrinsic behaviorally relevant neural dynamics and dissociate them from both other intrinsic dynamics and measured input dynamics. In data from a simulated brain with fixed intrinsic dynamics that performs different tasks, the method correctly finds the same intrinsic dynamics regardless of task while other methods can be influenced by the change in task. In neural datasets from three subjects performing two different motor tasks with task instruction sensory inputs, the method reveals low-dimensional intrinsic neural dynamics that are missed by other methods and are more predictive of behavior and/or neural activity. The method also uniquely finds that the intrinsic behaviorally relevant neural dynamics are largely similar across the three subjects and two tasks whereas the overall neural dynamics are not. These input-driven dynamical models of neural-behavioral data can uncover intrinsic dynamics that may otherwise be missed.

## Introduction

Neural population activity exhibits rich temporal structures^1–26^. Investigating these temporal structures, i.e., dynamics, can reveal the neural computations that underlie behavior^5,6,12,15,16,19,20^. Much progress has been made in developing models that can describe the dynamics of neural population activity using a low-dimensional latent state^2–4,7,8,10–14,16^. However, a major challenge in such investigations is that neural dynamics can arise due to two distinct sources that reflect distinct computations^12,15,27^. The first source consists of the intrinsic dynamics within a given brain region. Intrinsic dynamics arise due to the recurrent connections within a region’s neuronal population as it responds in a temporally structured manner to any excitations from within or outside that region^6,12,15,18,27,28^. The second source consists of input dynamics. These input dynamics are temporal structures that already exist in inputs that reach the recorded brain region, including sensory inputs or inputs from other brain regions^1,9,12,15,27,29–31^. While measuring all inputs is infeasible experimentally, measurements of sensory inputs such as task instructions or partial measurements of neural inputs into a brain region are often possible. As such, correctly interpreting how neural computations in a given brain region give rise to a specific behavior can greatly benefit from simultaneously achieving two objectives, which remain elusive.

First, given the above two sources, neural dynamics that are intrinsic to a given brain region need to be dissociated from those that are simply due to temporally structured measured inputs to that region. Second, within intrinsic neural dynamics, those that are relevant to the specific behavior of interest need to be dissociated from other intrinsic neural dynamics. This latter dissociation is important because neural dynamics of a specific behavior often constitute a minority of the total variance in the recorded neural activity^5,6,19,32–39^. Consistent with this observation, recent work has shown that learning dynamical models of neural-behavioral data together and in a way that dissociates and prioritizes their shared dynamics can unmask behaviorally relevant neural dynamics that may otherwise not be found^19,20^. We refer to this prioritized learning approach for neural-behavioral data as preferential dynamical modeling because it preferentially models the behaviorally relevant neural dynamics with priority instead of non-preferentially modeling prevalent dynamics in neural data as is typically done. But prior methods for preferential dynamical modeling cannot account for the effect of measured inputs to a given brain region^19^. Thus, the dissociation of intrinsic and input-driven neural population dynamics that underlie specific behaviors has remained challenging.

Here, we first show how misinterpretation and incorrect identification of intrinsic behaviorally relevant dynamics could result from modeling neural activity while considering behavior but not input, or while considering input but not behavior. Indeed, modeling neural activity without considering the measured input could result in a model that mistakes the temporal structure in the input as part of the intrinsic dynamics within the recorded brain region^9,27^ and consequently confound scientific conclusions. For non-preferential modeling of neural activity on its own, while not commonly done, various methods can be adapted to fit models with measured inputs^40^, thus accounting for measured input and neural activity but not behavior. However, as we show, even these non-preferential methods with input can miss those intrinsic neural dynamics that are behaviorally relevant. Further, methods for preferential modeling that consider the neural-behavioral data together cannot consider measured inputs. These results motivate the critical need for developing novel methods that can simultaneously consider neural activity, behavior, and measured inputs when learning a dynamical model (see Discussion). It is also important to recognize that disentanglement of intrinsic and input dynamics is fundamentally limited by the extent to which measured inputs are available, which depends on the experimental capability for input measurement. Perfect disentanglement requires measuring all inputs, which is typically not feasible with current experimental technology. As such, our aim here is to mathematically formulate a learning problem that involves neural activity, behavior, and measured inputs simultaneously, and to dissociate the intrinsic behaviorally relevant neural dynamics from the dynamics of any measured inputs and from other intrinsic neural dynamics. As we will show in our results, even this partial dissociation using the measured inputs (e.g., task instructions) can already lead to more accurate models and inferences, and to useful new insights compared with prior methods that account for either measured input or behavior during learning but not both.

With the above motivation, we then provide a new preferential modeling approach, termed Input Preferential Subspace Identification (IPSID) that can consider both measured inputs and behaviors in the training set while learning dynamical models of neural population activity. By doing so, IPSID provides the new capability to learn the intrinsic behaviorally relevant neural dynamics *with priority* and dissociate them both from other intrinsic neural dynamics and from the dynamics of measured inputs. We also develop a version of IPSID that achieves this capability when some input dynamics influence the behavior through pathways that are neither recorded nor downstream of the recorded neural activity. Compared with prior preferential modeling methods (i.e., PSID)^19,41^, which cannot incorporate input and thus do not dissociate intrinsic and input dynamics, IPSID requires distinct mathematical operations including completely new steps (**Note S1**). We show that two new capabilities provided by IPSID are critical for accurate dissociation of intrinsic behaviorally relevant neural dynamics: prioritized learning of these dynamics in the presence of input, and ensuring all learned dynamics are directly present in the neural recordings even when inputs affect behavior.

We validate IPSID and its new capabilities in extensive numerical simulations of diverse dynamical systems and in two independent motor cortical datasets from three non-human primates (NHP) recorded during two different tasks with task instruction sensory inputs. First, we simulate a brain with fixed intrinsic dynamics that performs different behavioral tasks. IPSID correctly learns the same intrinsic behaviorally relevant neural dynamics regardless of which specific task is used to collect the simulated training neural data. In contrast, other methods learn intrinsic dynamics that are inaccurate and influenced by the specific task. Second, we apply IPSID to motor cortical population activity recorded from three NHPs in two independent datasets with two different 2-dimensional (2D) cursor-control tasks. IPSID finds intrinsic behaviorally relevant dynamics that not only predict motor behavior better than non-preferential methods even with input, but also predict neural activity better than preferential methods which cannot consider task instruction inputs. Further, IPSID reveals that intrinsic behaviorally relevant neural dynamics are largely similar across the three animals despite differences in the two cursor-control tasks and animals, while other methods miss these similar dynamics. By dissociating intrinsic behaviorally relevant dynamics from both other intrinsic dynamics and measured input dynamics, IPSID can help explore unanswered questions regarding how intrinsic and input-driven neural computations give rise to behavior across subjects and tasks.

## Methods

### Modeling intrinsic neural dynamics underlying behavior in the presence of inputs

To see the effect of input on misinterpretation of intrinsic neural dynamics, consider a task where a subject is instructed to follow an on-screen target with their hand while motor cortical activity that represents the hand movements is recorded (**Fig. 1a**). Here, movements of the target would result in corresponding movements in the hand that follows the target, and thus would also introduce corresponding dynamics in the neural activity that represents hand movements. Consequently, any arbitrary movement of the target will be, to some extent, reflected in the recorded neural activity. An example is shown in a numerical simulation in **Fig. 1a,b**. As another example, if the target moves up and down with a 1s period, one would expect the neural activity to also include similar periodic patterns with a 1s period. If the period of target movements changes to 2s, so would the period of the patterns in neural activity that represent the hand movements. Any neural modeling that is not informed by target movements, which serve as task instruction sensory inputs, cannot distinguish between such input dynamics and intrinsic dynamics that originate in the recorded brain region. Thus, such modeling may incorrectly conclude that there exist intrinsic dynamics originating in the recorded brain area that are periodic with a 1s period. This means that sensory inputs or inputs from other brain regions, if unaccounted for, may confound dynamical models of neural activity by being misinterpreted as intrinsic dynamics. The reflection of input dynamics in neural dynamics can also be seen in terms of the frequency domain spectrum of these signals (**Fig. 1b**). In this view, the correct dissociation of intrinsic dynamics from input dynamics requires the correct learning of the transfer function from inputs to neural signals, in a way that doesn’t incorrectly attribute the input dynamics that appear in neural activity to having originated from the transfer function (**Fig. 1b**).

**Fig. 1.**
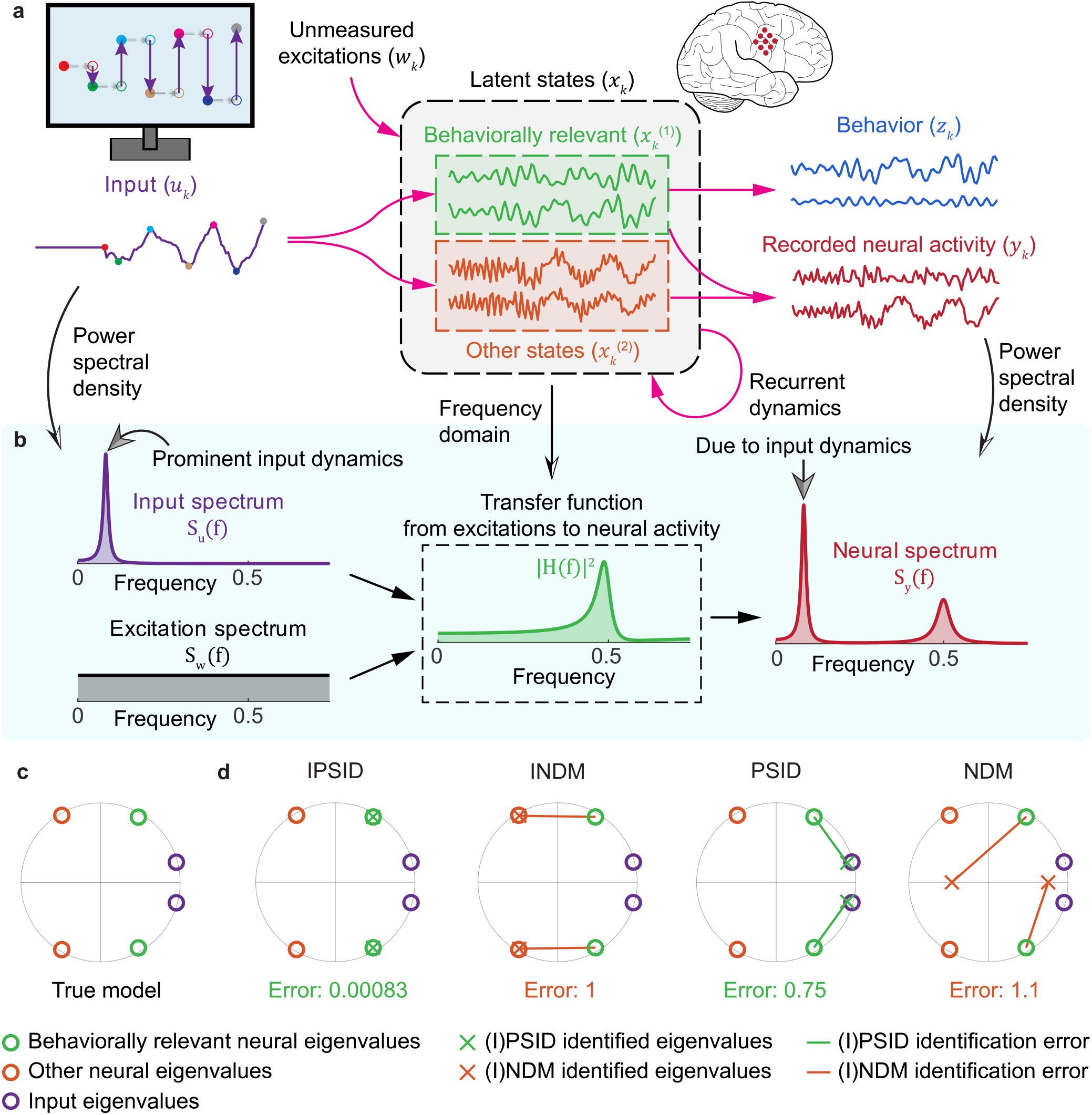
Intrinsic behaviorally relevant neural dynamics may be confounded by other intrinsic neural dynamics as well as by measured input dynamics, a challenge that the new IPSID method resolves. **(a)** Data generated from a simulated brain following equation (1) with a 1D input and a 4D latent state out of which only 2 dimensions (green) drive behavior. The input is taken as the sensory input such as target position moving up and down on a screen as depicted, but input can also consist of measured activity from other upstream brain regions. Neural dynamics that arise from the recurrent dynamics of neuronal networks within the brain region constitute the intrinsic neural dynamics. Oscillating temporal pattens of the input (left) constitute the input dynamics and clearly also appear in the neural activity (right) in a way that is mixed with the intrinsic neural dynamics. **(b)** Appearance of input dynamics in neural dynamics can also be clearly seen in the frequency domain representation of (a), showing: the power spectral density (PSD), or spectrum, of input time series *S_u_*(*f*) (top-left); PSD of unmeasured excitations *S_w_*(*f*) modeled as white Gaussian noise (bottom-left); transfer function from inputs to the neural activity (middle); and PSD of neural activity (right). Neural activity exhibits two dominant frequency components. In this simulation, the lower-frequency component is the reflection of input dynamics while the higher-frequency component represents intrinsic neural dynamics (as it is also present in the transfer function). Horizontal axes show the normalized frequency with 1 being the maximum possible frequency, i.e., *π*. **(c)** The eigenvalues of the latent state transition matrix *A* in the simulated brain model in equation (1). **(d)** Learned eigenvalues using (I)PSID or (I)NDM. Red lines indicate the error in the learned eigenvalues. The normalized error value—average line length normalized by the average true eigenvalue magnitude—is noted below each plot. Only IPSID correctly learns the intrinsic behaviorally relevant neural dynamics as quantified by the eigenvalues (**SI Methods**). Unlike IPSID, NDM or PSID may not learn the correct intrinsic dynamics but instead learn dynamics (eigenvalues) that are deflected towards the input dynamics (eigenvalues) in this example.

When performing non-preferential dynamic modeling of neural activity on its own, though not common, various methods such as subspace identification^40^ can be leveraged to fit models with measured input. However, as we will show, these methods can lead to inaccurate identification of intrinsic behaviorally relevant neural dynamics as behavior is not considered during learning. Further, current methods cannot account for measured inputs in preferential modeling of neural-behavioral data together, which we now enable by developing IPSID. To formulate the goal of IPSID, we represent the dynamical state of the recorded brain regions as a high-dimensional vector, of which each dimension may or may not contribute to generating the specific behavior of interest, i.e., be behaviorally relevant (**Fig. 1a**). Thus, in investigations concerned with behavior in the presence of a measured input, there are two major confounding factors in learning intrinsic behaviorally relevant neural dynamics: (1) the dynamics of the measured input that may be incorrectly considered part of intrinsic behaviorally relevant neural dynamics, and (2) other intrinsic neural dynamics that may mask or confound the behaviorally relevant ones.

IPSID addresses both confounding factors by accounting for neural activity, behavior, and measured input simultaneously during learning via new algebraic operations. Doing so, IPSID models and dissociates the intrinsic behaviorally relevant neural dynamics from measured input dynamics and from other intrinsic neural dynamics. Unlike IPSID, prior methods address only one or the other confound but not both. First, non-preferential neural dynamic modeling (NDM) with input (**SI Methods**), which we term INDM, accounts for neural activity and measured input but not behavior during learning. As such, INDM may miss or confound the intrinsic neural dynamics that are behaviorally relevant. Second, a method termed PSID^19,41^ addresses the second confound by accounting for neural activity and behavior during learning but not input. As such, PSID cannot dissociate intrinsic and input dynamics. We thus use this naming convention for ease of exposition but the algebraic operations in IPSID are different from those in both PSID and INDM, and further IPSID requires additional new mathematical steps compared with these prior methods (**Notes S1, S2**).

In IPSID, we use the following linear state-space model to jointly describe the dynamics of neural activity (*y_k_*) and behavior (*z_k_*) in the presence of measured input (*u_k_*)

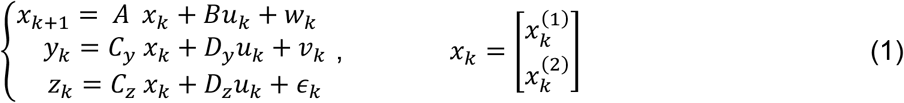

where 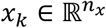 is the latent neural state composed of two parts: (1) 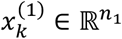, which is the behaviorally relevant latent states in the recorded neural activity; (2) 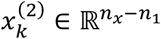, which is the other latent states in the recorded neural activity. In this model, 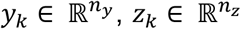 and 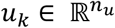 represent the recorded neural activity, the measured behavior, and the measured input, respectively. Here 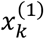 being behaviorally relevant means that only those dimensions of *x_k_* corresponding to 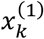 contribute to generating behavior (*z_k_*) in the third row of equation (1). Finally, *w_k_* and *v_k_* are zero mean white Gaussian noises (**SI Methods**) and *ϵ_k_* is a general Gaussian random process representing any behavior dynamics not encoded in the recorded neural activity (i.e., not driven by *x_k_*).

Prior works have not addressed the problem of fitting this model in a way that dissociates and prioritizes the learning of behaviorally relevant latent states. To enable such preferential/prioritized learning, we developed IPSID, which uses new linear algebraic operations to directly extract the subspace of intrinsic behaviorally relevant latent states from neural, behavioral and input training data, and then learns the model parameters. IPSID provides a new two-stage learning procedure that incorporates input as follows. In the first stage of IPSID, we develop algebraic operations that extract the behaviorally relevant latent states with *priority* via an oblique (non-orthogonal) projection of future behavior onto past neural activity and past inputs along the subspace spanned by future inputs (**Fig. S1, SI Methods**). Then, in an optional second stage, we devise algebraic operations that extract any other latent neural states by another oblique projection from any residual/unexplained future neural activity onto past neural activity and past inputs along future inputs (**Fig. S1**). Model parameters are then learned via least squares based on the extracted latent states and their relation in equation (1). This two-stage learning enables the new capability for prioritized learning of the intrinsic behaviorally relevant neural dynamics over other intrinsic neural dynamics in the presence of inputs because the former dynamics are learned first, i.e., in the first stage. The two-stage learning allows us to learn a minimally complex model of the intrinsic behaviorally relevant neural dynamics in the first stage (i.e., a model with low-dimensional states), instead of having to learn a more complex model that simultaneously includes all intrinsic neural dynamics. This leads to learning more accurate models of intrinsic behaviorally relevant dynamics for a given dataset as shown in extensive simulations and in real neural data analyses below. After the model is learned, in the test set, extraction of intrinsic behaviorally relevant neural dynamics is done without looking at behavior and via a Kalman filter associated with the learned model (**SI Methods**). Details of IPSID are provided in **SI Methods** and **Notes S1–S2**.

We found that among the learning methods, the only method that correctly learns the intrinsic behaviorally relevant neural dynamics in the presence of inputs is the new IPSID (**Fig. 1d**) as demonstrated below. Further, by implementing a numerical optimization approach that maximizes the data likelihood with block-structured constraints on model parameters (**SI Methods**), we demonstrate the critical importance of two new capabilities provided by IPSID for dissociating the intrinsic behaviorally relevant neural dynamics: i) *prioritized* learning of these dynamics enabled via the new two-stage learning algorithm in the presence of input as described above, and ii) ensuring all learned latent states are directly present in neural recordings even when inputs affect behavior. To assess the methods, we look at the eigenvalues of the latent state transition matrix *A*, which quantify the dynamics (**SI Methods**, **Fig. 1c,d**). We also compute the accuracy in decoding behavior from neural activity as well as in neural self-prediction – defined as predicting neural activity one-step-ahead from its own past (**SI Methods**).

## Results

### IPSID correctly learns all model parameters in the presence of inputs

We first validated the accurate learning of intrinsic behaviorally relevant neural dynamics using IPSID in a simulated model with 4-dimensional latent states out of which only two dimensions were involved in generating the simulated behavior (**Fig. 1a**). The eigenvalues of the state transition matrix *A* affect the transfer function from the input to the states and neural activity (**Fig. 1b**), characterize the state response to excitations, and describe the dynamics (**Fig. 1c**, **SI Methods**). We thus use these eigenvalues to quantify the intrinsic neural dynamics (**SI Methods**). We found that the new IPSID was the only method that correctly learned the eigenvalues associated with the intrinsic behaviorally relevant neural dynamics (**Fig. 1d**). In contrast, NDM or PSID that do not consider inputs learned models that were confounded by input dynamics and INDM that does not consider behavior was confounded by other intrinsic neural dynamics beyond the behaviorally relevant ones.

To more generally validate IPSID, we applied it to data generated from 100 random models in the form of equation (1) with random parameters and dimensions (**SI Methods**). To provide input to these models, we independently simulated another 100 models without input (equation (3) from **SI Methods**) with random parameters and passed their output as the input to the original models—these inputs are thus generated by an independent dynamical system and can be thought of as activity of other brain regions or as structured sensory inputs. We found that IPSID correctly learned all model parameters in the presence of inputs (**Fig. S2**). Moreover, the rate of convergence of parameters as a function of training samples was similar to INDM (**Fig. S2b**); this suggests that despite its additional capability in dissociating those intrinsic neural dynamics that are behaviorally relevant, IPSID does not require more training data than INDM even when modeling all latent states.

### IPSID correctly prioritizes the learning of intrinsic behaviorally relevant neural dynamics in the presence of inputs

In another numerical simulation, we found that IPSID correctly prioritizes the learning of intrinsic behaviorally relevant neural dynamics in the presence of inputs, which is an important new capability for learning these dynamics accurately (**Fig. 2,** also see **Fig. 4** later). Thus, IPSID addresses the challenge of preferential dynamical modeling of neural-behavioral data with inputs (**Fig. 2**). We simulated 100 random models formulated by equation (1) with a 6D latent state, out of which only 2 dimensions were behaviorally relevant (**SI Methods**). To get the input to these models, we independently simulated 100 random models without input (equation (3) from **SI Methods**) with 2D latent states and passed their output as the input to the original models. We then learned models using (I)PSID and (I)NDM with varying latent state dimensions (*n_x_*). In each case, we computed the error in learning the intrinsic behaviorally relevant eigenvalues, which quantifies how accurately intrinsic behaviorally relevant dynamics are learned (**Fig. 2b**, **Fig. S3**).

**Fig. 2.**
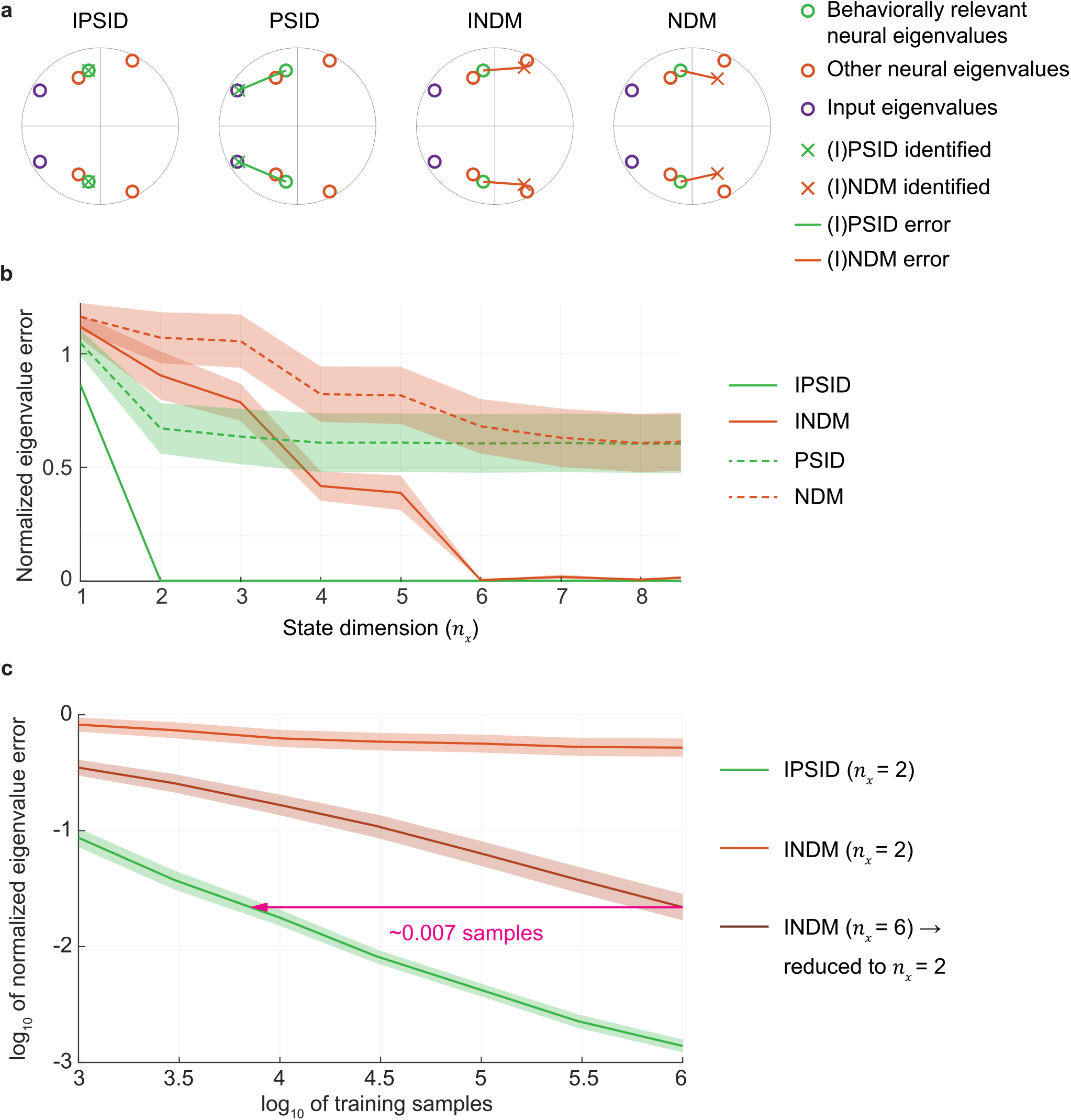
IPSID correctly prioritizes the learning of intrinsic behaviorally relevant neural dynamics thus achieving preferential neural-behavioral modeling even in the presence of input. **(a)** For one simulated model (equation (1)), the identified intrinsic behaviorally relevant eigenvalues are shown for (I)PSID and (I)NDM using a 2D latent state. Eigenvalues of the state transition matrix *A* in the true model are shown as colored circles. Crosses show the identified behaviorally relevant eigenvalues when modeling the neural activity. **(b)** Normalized error of learning the intrinsic behaviorally relevant eigenvalues given 10^6^ training samples is shown when using (I)PSID and (I)NDM, averaged over 100 random models each with total latent state dimension of *n_x_* = 6 and behaviorally relevant state dimension of *n*_1_ = 2. For all models, an independent random model with state dimension of 2 generated the input (**SI Methods**). Solid lines show the average across models and shaded areas show the s.e.m. (*n* = 100 random models). For all methods, we vary the state dimension *n_x_* from 1 to 8; for *n_x_* > 2, we find the 2 state dimensions that best predict behavior and evaluate their 2 associated eigenvalues (**SI Methods**). We find that to learn the intrinsic behaviorally relevant eigenvalues, IPSID only needs a minimal state dimension *n_x_* = 2 (true *n*_1_) whereas INDM needs a high state dimension *n_x_* = 6 (true total model dimension *n_x_*). This also leads to INDM’s higher error with the same training sample size in (c). Also, even using high-dimensional states, NDM and PSID cannot dissociate which of the eigenvalues are intrinsic and thus do not learn the correct reduced models (because they do not consider input). **(c)** Normalized error of learning the intrinsic behaviorally relevant eigenvalues vs. training samples for 100 random models. For INDM, we try i) directly learning a model with a 2D latent state and ii) first learning a model with a high enough dimension to achieve almost zero error in (b) and then reducing the model to keep the top 2 dimensions with the best behavior decoding (indicated by dimension → 2) (**SI Methods**). INDM requires orders of magnitude more training samples than IPSID to learn the intrinsic behaviorally relevant eigenvalues with similar accuracy.

**Fig. 3.**
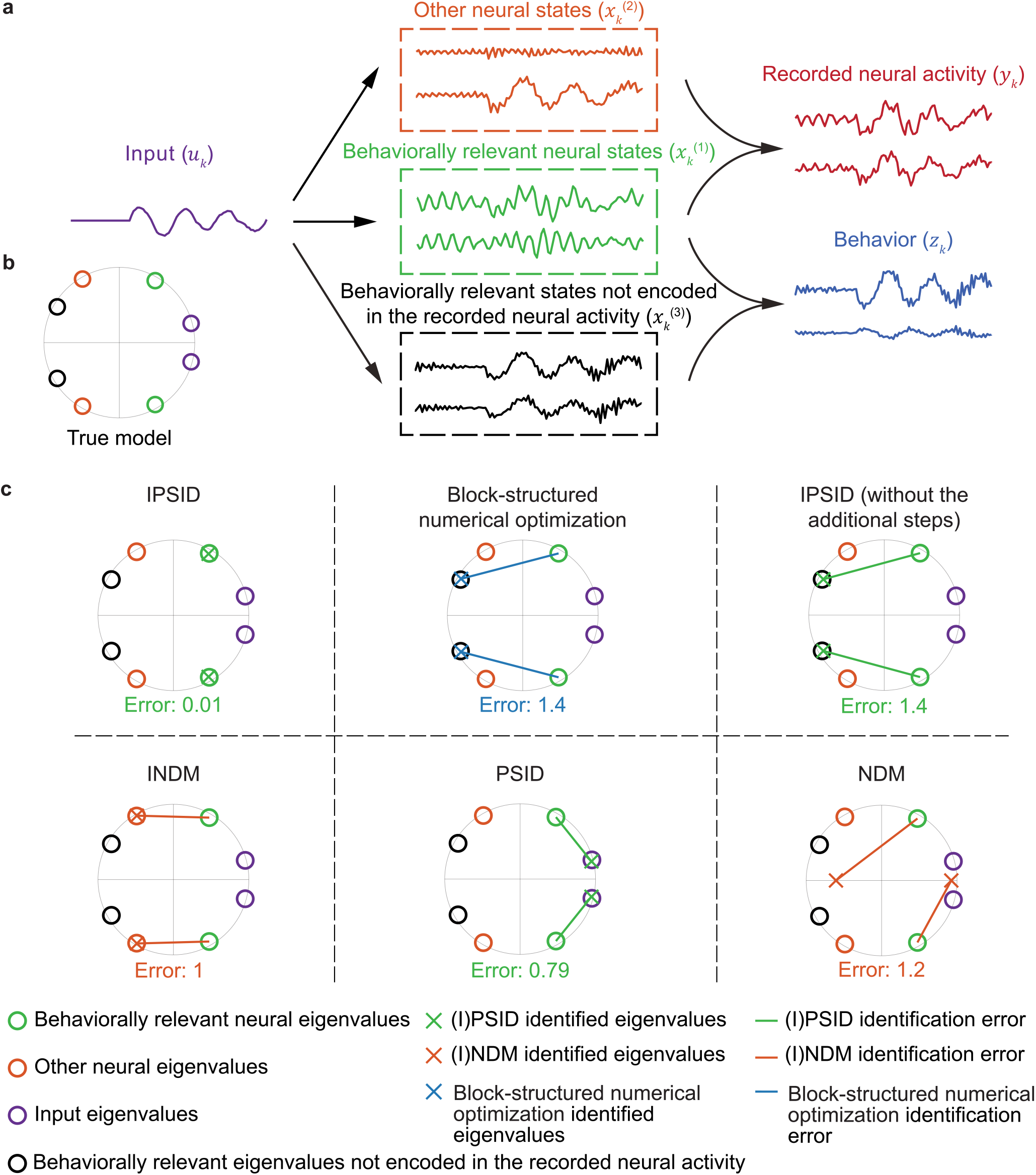
IPSID also applies to scenarios when the recorded regions do not cover all downstream regions of the input. **(a)** A simulated brain (as in equation (2)) with a 6D latent state out of which only 4 dimensions drive the recorded neural activity and the other 2 dimensions just drive the behavior. **(b)** The eigenvalues of the state transition matrix *A* in the simulated model. The 4 eigenvalues associated with the 4 state dimensions that drive the recorded neural activity are shown as green and orange circles, depending on whether they drive behavior (green) or not (orange). Eigenvalues associated with the two additional state dimensions that only drive the behavior but not recorded neural activity are shown as black circles. **(c)** Eigenvalues of the models learned using IPSID, block-structured numerical optimization, IPSID (without additional steps), PSID and (I)NDM. A simplified schematic of key operations for each method is in **Fig. S6**. The block-structured numerical optimization learns the model parameters via gradient descent (**SI Methods**). Notation is as in **Fig. 1**. IPSID can also address this scenario using its additional steps. Only IPSID correctly learns the intrinsic behaviorally relevant neural dynamics even in this scenario, and its new capability via the additional steps is needed to avoid the black eigenvalues/dynamics (behaviorally relevant dynamics not reflected in the recordings; see the top row comparisons).

**Fig. 4.**
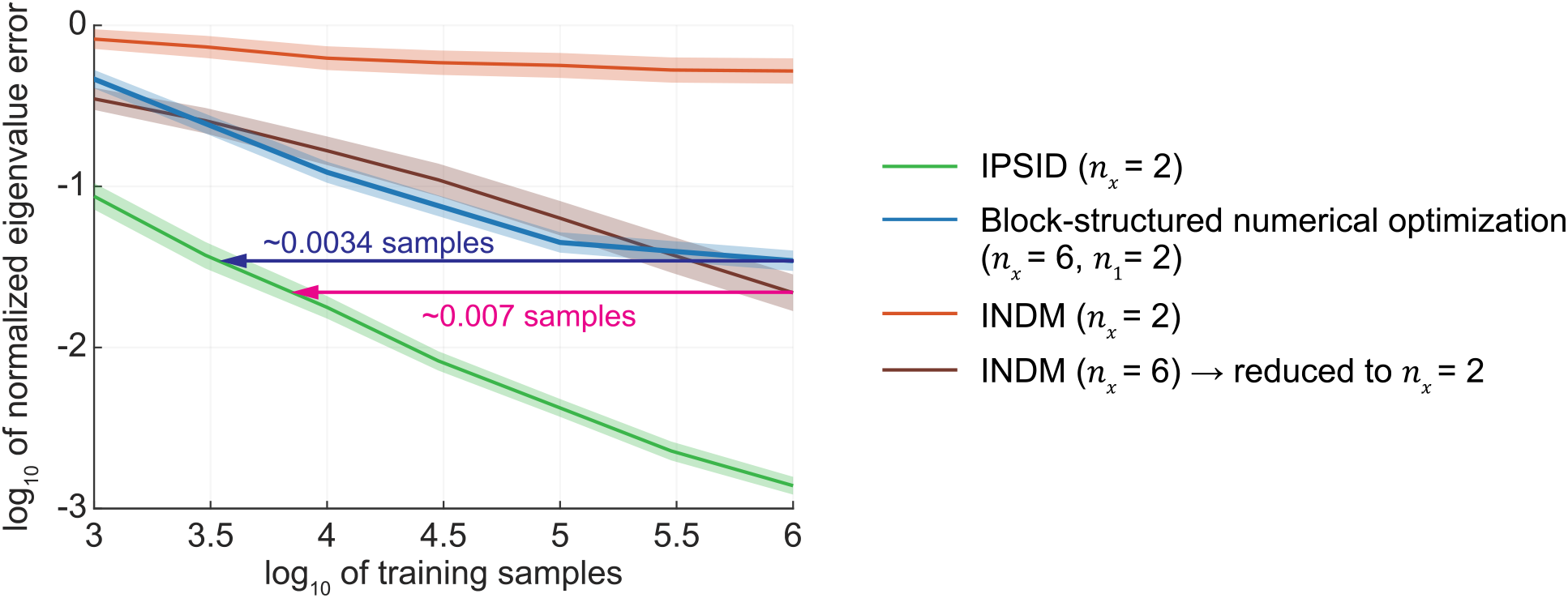
IPSID outperforms numerical optimization with block-structured model parameters in terms of accuracy for model learning, showing the importance of prioritized learning and dissociation. Notation is as in **Fig. 2c**, but also shows the error of learning the intrinsic behaviorally relevant eigenvalues for an additional learning approach based on numerical optimization with block-structured model parameters (**SI Methods**). Similar to INDM (which is replicated from **Fig. 2c** here), this optimization approach is significantly less accurate for a given training data (i.e., number of samples). Also, it requires orders of magnitude more training samples than IPSID to learn the intrinsic behaviorally relevant eigenvalues as accurately.

We found that only IPSID could learn all the intrinsic behaviorally relevant neural dynamics using the minimal latent state dimension of 2, which is the true simulated dimension of these dynamics (**Fig. 2b**, **Fig. S4**). IPSID was able to achieve this by considering both inputs and behavior in its preferential modeling of neural dynamics during learning. This meant that IPSID could simultaneously dissociate the intrinsic behaviorally relevant neural dynamics form other intrinsic neural dynamics and from input dynamics. In contrast, NDM and PSID do not consider the input and thus were unable to dissociate the intrinsic versus input dynamics, leading to a high eigenvalue error (**Fig. 2b**). Further, even though INDM considers inputs, it does not consider behavior during learning and thus it required much larger latent state dimensions to learn the intrinsic behaviorally relevant eigenvalues (**Fig. 2b**). Also, even INDM with a higher state dimension (i.e., 6) had larger eigenvalue error when using the same number of training samples as IPSID (**Fig. 2c**); this is because models with higher dimensional states are more complex and thus more difficult to learn. Indeed, IPSID required orders of magnitude fewer training samples to learn the intrinsic behaviorally relevant neural dynamics in the presence of inputs (**Fig. 2c**).

We next found that NDM and PSID models, which do not consider input, could not accurately learn the intrinsic behaviorally relevant dynamics even by increasing their state dimension. Specifically, we first learned an NDM/PSID model with a high latent state dimension that learns a mixture of all intrinsic neural dynamics and input dynamics. We then reduced this model, as we did with INDM above, by only keeping the two dimensions that were best in decoding behavior and looking at their associated eigenvalues (**SI Methods**). Even with this approach, the reduced models were still much less accurate than low-dimensional models learned with IPSID (see **Fig. 2b** at high dimensions).

### IPSID can dissociate the effects of input on behavior that are reflected in the recorded neural activity from those that are not

In equation (1), all the effects of input on behavior happen through latent states that are reflected in the recorded neural activity. In this scenario, all the downstream regions of the input are either covered in the recordings or reflected in them (e.g., are downstream of the recorded regions). In addition to this scenario, we now show that IPSID can also apply to a new scenario where inputs may also influence behavior through pathways/regions that are neither recorded nor reflected in the recorded activity (**Fig. 3a**). We formulate this new scenario with the following model

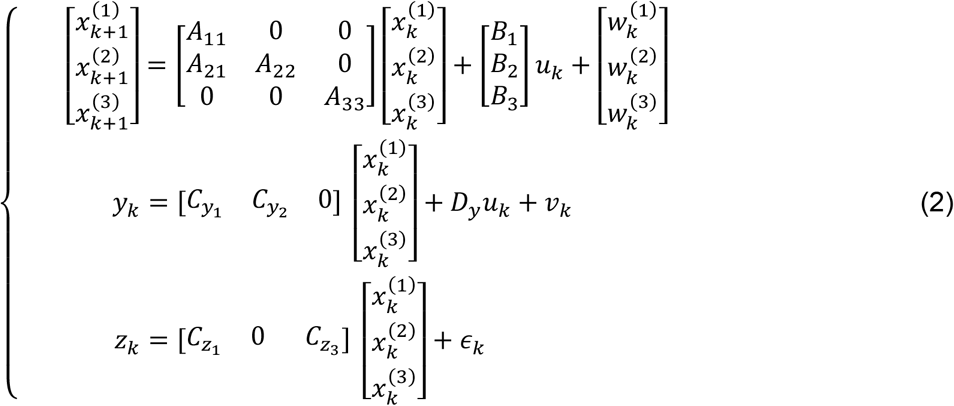

where compared with equation (1), an additional segment 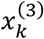 is added to the latent state *x_k_* to represent the effects of input on behavior *z_k_* that are not reflected in the recorded neural activity *y_k_*. In this formulation, IPSID dissociates the latent state into three segments: (1) 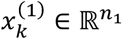, which is the behaviorally relevant latent state that is reflected in neural activity *y_k_*, (2) 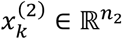, which is the latent state that describes any other neural dynamics, and (3) 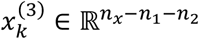, which is the behaviorally relevant latent state not reflected in the recorded neural activity *y_k_*. These three types of latent states are shown in an example in **Fig. 3a**. Note that in this case, only 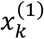 and 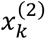 constitute the intrinsic latent states because only these latent states drive the recorded neural activity. To add support for dissociation of these three types of latent states to IPSID, we developed two additional optional steps for IPSID (**Fig. S5**, **Note S2**).

In the first additional step, before the initial oblique projection of behavior onto neural activity and input, we project behavior onto the subspace of latent states in neural activity (i.e., neural states) irrespective of the relevance of these states to behavior; these neural states are obtained using only the second stage of IPSID (**SI Methods**, **Note S2**, **Figs. S5, S6a**). We then apply IPSID as before (**Note S1**), but now use the results of this additional projection as the behavior signal. This additional projection ensures that behavior dynamics that are not encoded in the recorded neural activity are not included in the first set of states 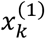.

In the second additional step, we optionally extract 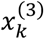, which represents any behavior dynamics that are driven by the input but are not encoded in the recorded neural activity. This means that 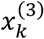 reflects processing in the downstream regions of input that are not recorded/reflected as part of neural activity. In this step, after performing the first additional step above and subsequently both stages of IPSID to extract 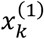 and 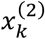, we compute the residual behavior that is still not predictable using 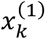 and 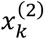. Then, using the second stage of IPSID, we build a model that predicts these residual behavior dynamics purely using the input (**SI Methods**, **Note S2**, **Fig. S5**) – this gives 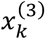, which summarizes the direct effect of input on behavior dynamics that are not reflected on the recorded neural activity. Together, these two additional steps enable IPSID to learn a model as in equation (2). If extraction of 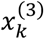 is not of interest, the second step can be optionally skipped, and solely the first step can be added to IPSID to ensure that 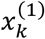 and 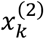 are encoded in the recorded neural activity.

We simulated models in the form of equation (2) and confirmed that with the above additional steps, IPSID correctly dissociates intrinsic behaviorally relevant neural dynamics (i.e., 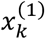) from other dynamics – i.e., from other intrinsic neural dynamics, input dynamics, and behavior dynamics not encoded in the recorded neural activity (**Fig. 3c**). Moreover, across 100 random models, IPSID correctly learned all model parameters in equation (2) (**Fig. S7**). Finally, by learning 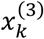, which captures the behavior dynamics that are predictable from input but are not reflected in the recorded neural activity, IPSID also achieved ideal prediction of behavior from input and neural activity (**Fig. S8**).

These results demonstrate that IPSID is applicable to scenarios where the recorded neural activity does not cover all the downstream regions of the measured input. In this scenario, IPSID can also dissociate the influences of input on behavior that are reflected in the recorded neural activity from those that are not. Without this capability, some of the learned dynamics may not be present in the recorded region (**Fig. 3c**, *top row comparisons*). Thus, this is another new capability by IPSID that is important for accurately dissociating intrinsic behaviorally relevant dynamics in neural recordings.

### Prioritized learning of intrinsic behaviorally relevant neural dynamics enabled with IPSID is critical for their accurate dissociation

A new capability provided by IPSID that is critical for disentangling intrinsic behaviorally relevant dynamics is to prioritize their learning over other intrinsic neural dynamics in the presence of input. This prioritized learning is enabled with IPSID’s new two-stage learning procedure that incorporates input. As shown earlier in **Fig. 2**, without prioritized learning, more latent states would be needed to ensure that intrinsic behaviorally relevant neural dynamics are included in the model, which will result in much higher error in learning these dynamics for a given training dataset (**Fig. 2c**). To further show the importance of the new prioritized learning capability, we implemented a block-structured numerical optimization approach that fits a model with the same block structure as the IPSID model in equation (6) from **SI Methods**. This optimization fits all model parameters to simultaneously maximize the neural-behavioral data log-likelihood (**SI Methods**). Note that the two-stage learning procedure in IPSID enforces a distinct learning objective, which is future behavior prediction in stage 1 and future residual neural prediction in stage 2. Thus, the IPSID objective is different from the numerical optimization objective, which is to simultaneously optimize the likelihood of neural and behavioral data. We applied the numerical optimization approach to the same simulated data as in **Fig. 2c**. We found that this block-structured numerical optimization approach is significantly less accurate than IPSID for a given number of training samples. Indeed, this approach requires orders of magnitude more training samples compared to IPSID to achieve the same accuracy (**Fig. 4** blue). This analysis shows that simply imposing a set of block-structured model parameters as in equation (6) is not sufficient for accurate disentanglement; rather the new capability for prioritized learning enabled by the two-staged learning approach in IPSID is important for achieving accurate disentanglement.

We also compared the computation times of the block-structured numerical optimization approach vs. IPSID. Model fitting using IPSID, which is based on a fixed set of linear algebraic operations, was significantly faster than model fitting using the numerical optimization approach (**Fig. S9**). Finally, this numerical optimization approach does not dissociate the effects of input on behavior that are reflected in the recorded neural activity from those that are not. Therefore, this approach may learn behavior dynamics not encoded in the recorded neural activity (**Fig. 3c**). This highlights the importance of the additional steps incorporated in IPSID as another new capability that is important for dissociating the intrinsic behaviorally relevant dynamics that are present in the recorded neural activity, as also explained in the previous section (**Fig. 3c, Fig. S5**, **Note S2**). Future work may be able to use similar ideas as the prioritized learning or these additional steps to develop numerical optimization approaches for the disentanglement problem formulated here (Discussion).

### Realistic motor task simulations show how sensory inputs to the brain can confound models of neural activity

As explained earlier (**Fig. 1a**), sensory inputs such as task instructions are effectively inputs to the brain that can have different dynamics from task to task, even if the intrinsic neural dynamics remain unchanged. Thus, unless accounted for during modeling, task-specific sensory inputs could confound the learned intrinsic neural dynamics. Developing a method that can learn the correct intrinsic neural dynamics regardless of the task would allow experimenters to study any given behavioral task of interest or compare intrinsic dynamics across different tasks without worrying about confounding the results and without limiting the task design. We hypothesized that even when the intrinsic neural dynamics remain unchanged, methods that do not consider the task sensory inputs may learn different and incorrect intrinsic dynamics depending on the exact task, whereas IPSID can learn the same intrinsic dynamics regardless of the task. Here we confirm this hypothesis by simulating a brain performing various realistic cursor control motor tasks during which simulated neural data for model training is observed (**Fig. 5**; **SI Methods**).

**Fig. 5.**
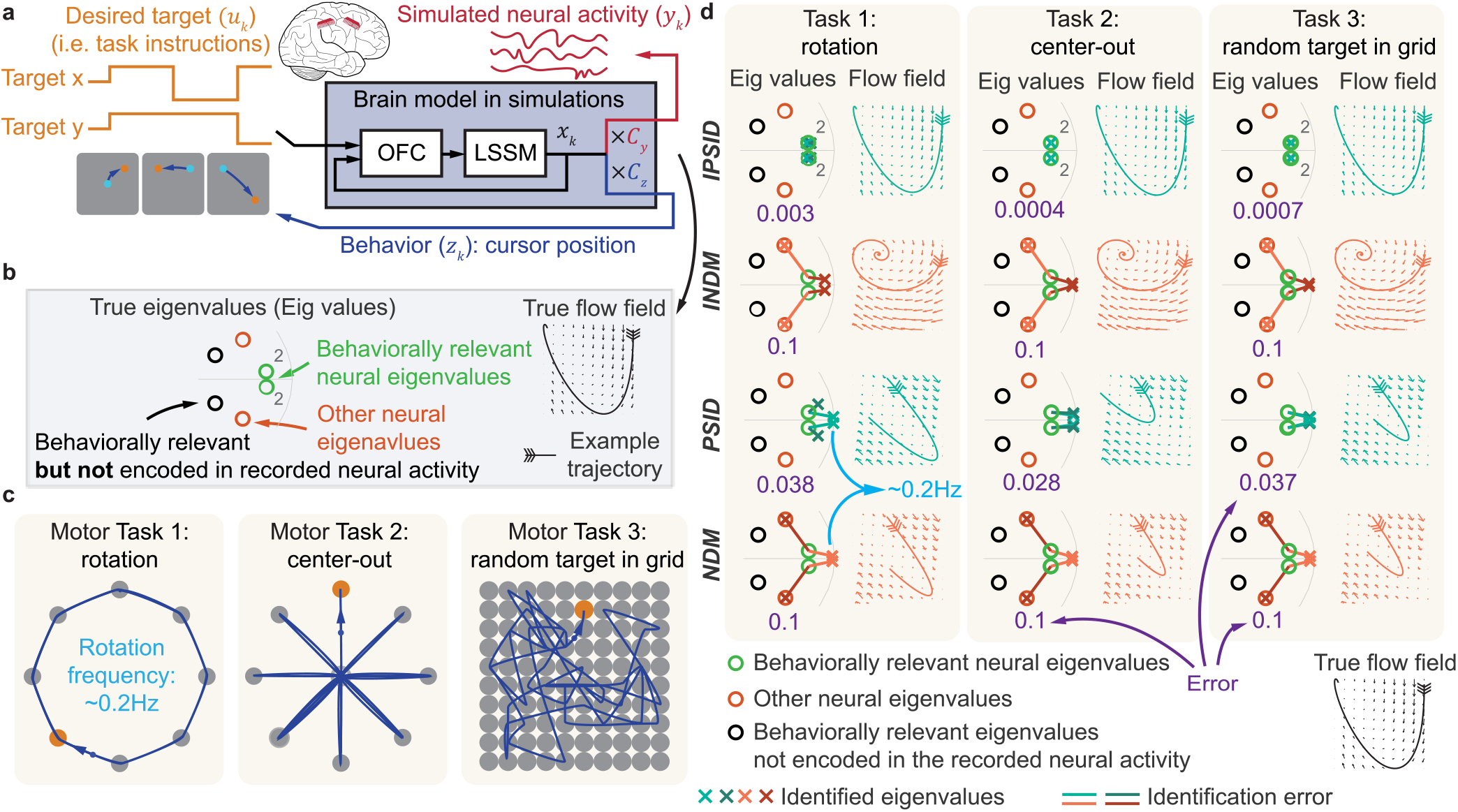
By considering task instruction inputs, IPSID learns the correct intrinsic behaviorally relevant neural dynamics regardless of the behavioral task unlike other methods. **(a)** The brain model consists of an optimal feedback controller (OFC) combined with a linear state space model (LSSM). Four of the 8 latent state dimensions of the LSSM encode the 2D position and velocity of the cursor (**SI Methods**). OFC controls these 4 state dimensions such that cursor position reaches the target shown on the screen while cursor velocity goes to zero (i.e., cursor stops at target). **(b)** Eigenvalues of the state transition matrix in the full brain model (i.e., OFC together with the LSSM) and the flow field associated with the behaviorally relevant neural eigenvalues. Flow fields show the direction in which the state would change starting from various initial values. In this brain model, there are two sets of behaviorally relevant complex conjugate eigenvalues that are in the same place and thus overlapping. Each set is associated with one movement direction, horizontal and vertical, respectively. The fact that there are two overlapping sets of eigenvalues is indicated by writing a 2 next to these eigenvalues. In addition to the 4 states representing position and velocity in the 2D space, there are 2 states that only drive the neural activity, whose associated eigenvalues are depicted as orange circles. There are also 2 states that only drive the behavior, whose associated eigenvalues are depicted as black circles. **(c)** Tasks performed by the simulated brain. **(d)** Identified eigenvalues for each task using each method with a state dimension of 4. The flow field for one of the two sets of eigenvalues identified by each method (the one with the lighter green/red color) is also shown as an example. Only IPSID correctly learns the intrinsic behaviorally relevant neural eigenvalues regardless of the behavioral task used during training.

Specifically, we modeled the brain as an optimal feedback controller^42–44^ (OFC), which controls a part of its state that represents the 2D cursor kinematics such that the cursor moves to targets presented via task instructions (**SI Methods**; **Fig. 5a**). The eigenvalues of the state transition matrix for the true simulated brain model are shown in **Fig.5b**. For generality, as part of the simulated brain, we included two latent states (similar to 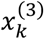 in equation (2)) that are driven by input and affect the movement (i.e. behavior), but are not reflected in the neural dynamics (**SI Methods**). As the first task, we simulated 8 equally spaced targets around a circle and instructed the simulated brain to move the cursor to the targets in order (**Fig. 5c**, left). As the second task, we simulated a standard center-out-and-back task where in each trial the cursor needs to move from the center to a randomly-specified target among 8 targets and then return back to the center (**Fig. 5c**, middle). Lastly, we simulated a 10 by 10 grid of targets where in each trial a random target within a limited distance of the most recent target needs to be visited (**Fig. 5c**, right) similar to the tasks in our NHP datasets (**SI Methods**). For each task, we used (I)PSID and (I)NDM to learn models of neural dynamics (**Fig. 5d**).

We found that regardless of the task, IPSID correctly learned the intrinsic behaviorally relevant neural dynamics. This is evident from comparing the IPSID eigenvalues and flow fields for every task with their ground-truth (first row of **Fig. 5d** vs. **Fig. 5b**). INDM, which considers input but not behavior during training, learned an approximation of some intrinsic behaviorally relevant neural dynamics with error, and also mistakenly included some intrinsic neural dynamics that were not relevant to behavior (**Fig. 5d**, second row). PSID, which considers behavior and neural activity but not input during training, learned biased intrinsic neural dynamics that were influenced by task instruction inputs (**Fig. 5d**, third row). Finally, NDM, which only considers neural activity during training, not only learned neural dynamics that were not related to behavior, but also learned inaccurate intrinsic behaviorally relevant neural dynamics that were influenced by task instruction inputs (**Fig. 5d**, fourth row). For example, in the first task, the biased dynamics learned by NDM and PSID were very close to the dominant frequency of the task instructions, which was around 0.2Hz (**Fig. 5d**, left column). These results demonstrate that by considering both behavior and sensory inputs such as task instructions during model training, IPSID can learn models of neural dynamics that are not confounded by the specific behavioral task during which neural data is collected. The ability to avoid these confounds is critical for comparing intrinsic neural dynamics across tasks in neuroscience investigations, as we also show in our real NHP neural data analyses below (**Fig. 8**).

**Fig. 6.**
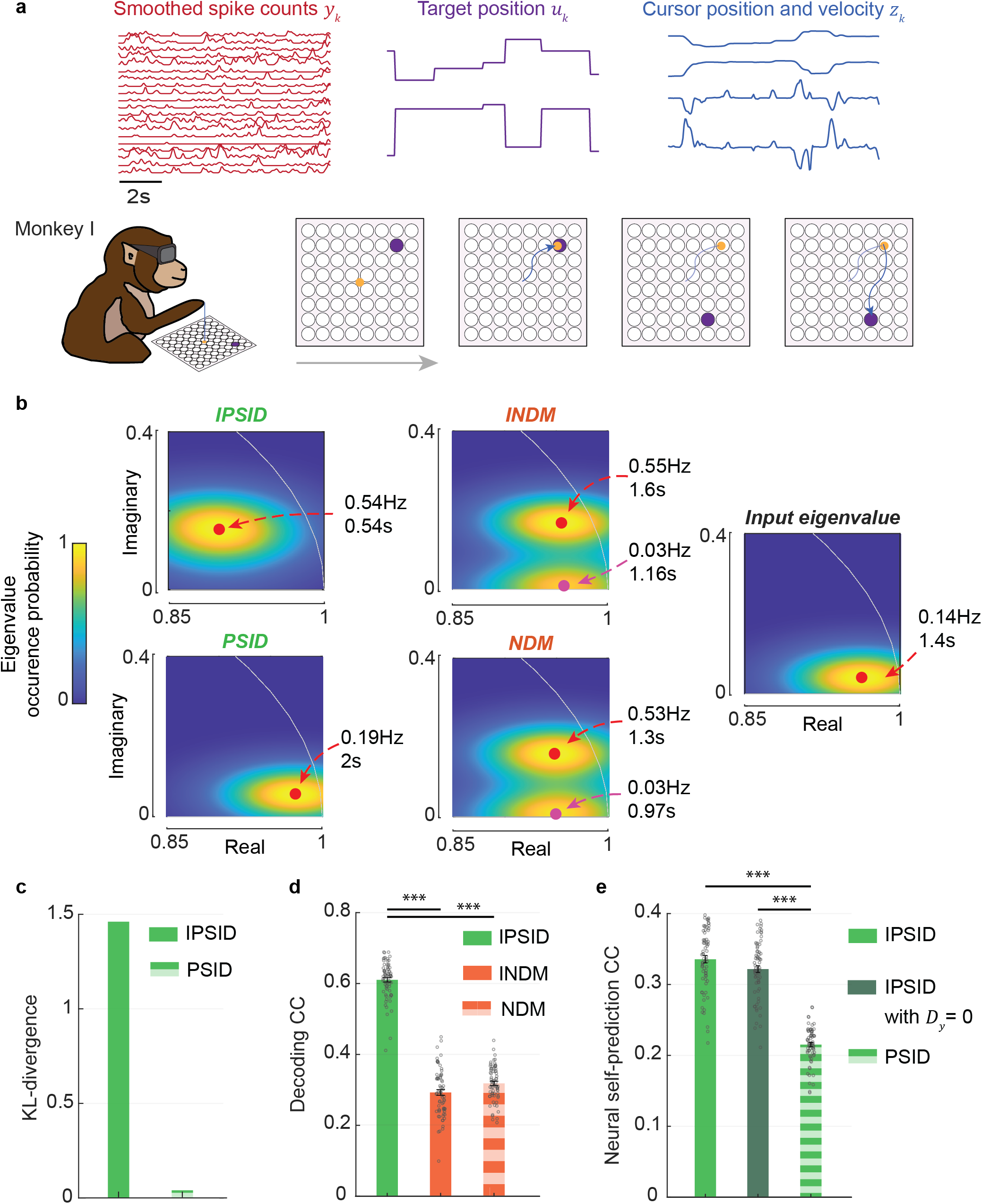
IPSID uncovers distinct and more accurate intrinsic behaviorally relevant neural dynamics in motor cortical population activity by considering task instructions as inputs to the brain. (**a**) We modeled the population spiking activity in a monkey (monkey I) performing a 2D cursor control task (**SI Methods**). See **Fig. S11** for results from a second monkey in this task and **Fig. 7** for results in a second dataset recorded from a different monkey in a different task. Spike counts are smoothed using a Gaussian kernel with s.d. of 50 ms (**SI Methods**). The 2D position and velocity of the cursor were taken as the behavior signal of interest and the target position time series was taken as the input to the brain. **(b)** Distribution of the eigenvalues of the state transition matrix for models learned using (I)PSID and (I)NDM across datasets. Input eigenvalue was found by applying NDM to the time-series of task instructions. Models were learned with a latent state dimension of *n_x_* = 4, which is sufficient for capturing most behavior dynamics (**Fig. S13**). We estimated the probability of an eigenvalue occurring at each location on the complex plane by adding Gaussian kernels centered at locations of all identified eigenvalues (*n* = 70 cross-validation folds across 2 channel subsets and 7 recording sessions, **SI Methods**). Red dots indicate the location that has the maximum estimated eigenvalue occurrence probability, with the associated frequency and decay rate (**SI Methods**) noted next to each plot. Similarly, when the occurrence probability map has more than one local maximum (i.e., for NDM or INDM), pink dots indicate the location of the second local maximum. **(c)** Quantified by KL-divergence, the eigenvalues learned by PSID were much closer to input eigenvalues than the eigenvalues learned by IPSID, showing the success of the new algebraic operations in accounting for inputs in neural-behavioral modeling. We computed the KL-divergence between the probability mass function of input eigenvalues (panel b, right) and the probability mass function of eigenvalues learned by IPSID/PSID (panel b, top/bottom left). **(d-e)** Cross-validated behavior decoding (panel d) and neural self-prediction (panel e) when modeling data with dimension *n_x_* = 4 and corresponding to models in (b). Triple asterisks indicate *P* < 0.0005 for a one-sided signed-rank test.

**Fig. 7.**
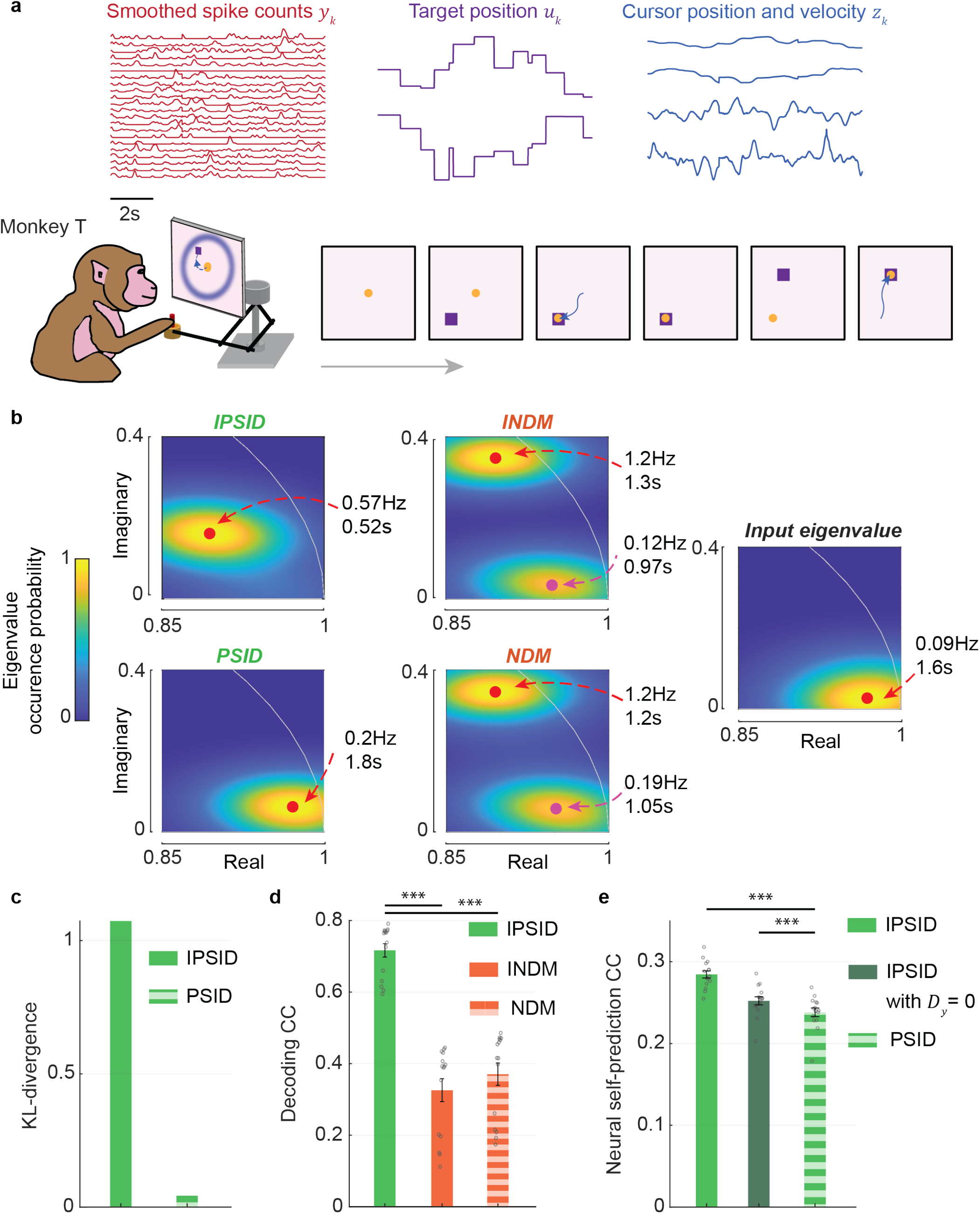
In a second dataset recorded from a different monkey and during a different task, IPSID again uncovers distinct and more accurate intrinsic behaviorally relevant neural dynamics in spiking activity by considering task instructions as inputs to the brain. Similar to **Fig. 6** for the second subject (monkey T, *n* = 15 cross-validation folds across 3 recording sessions, **SI Methods**) during a different second task with sequential reaches to random targets (**SI Methods**).

**Fig. 8.**
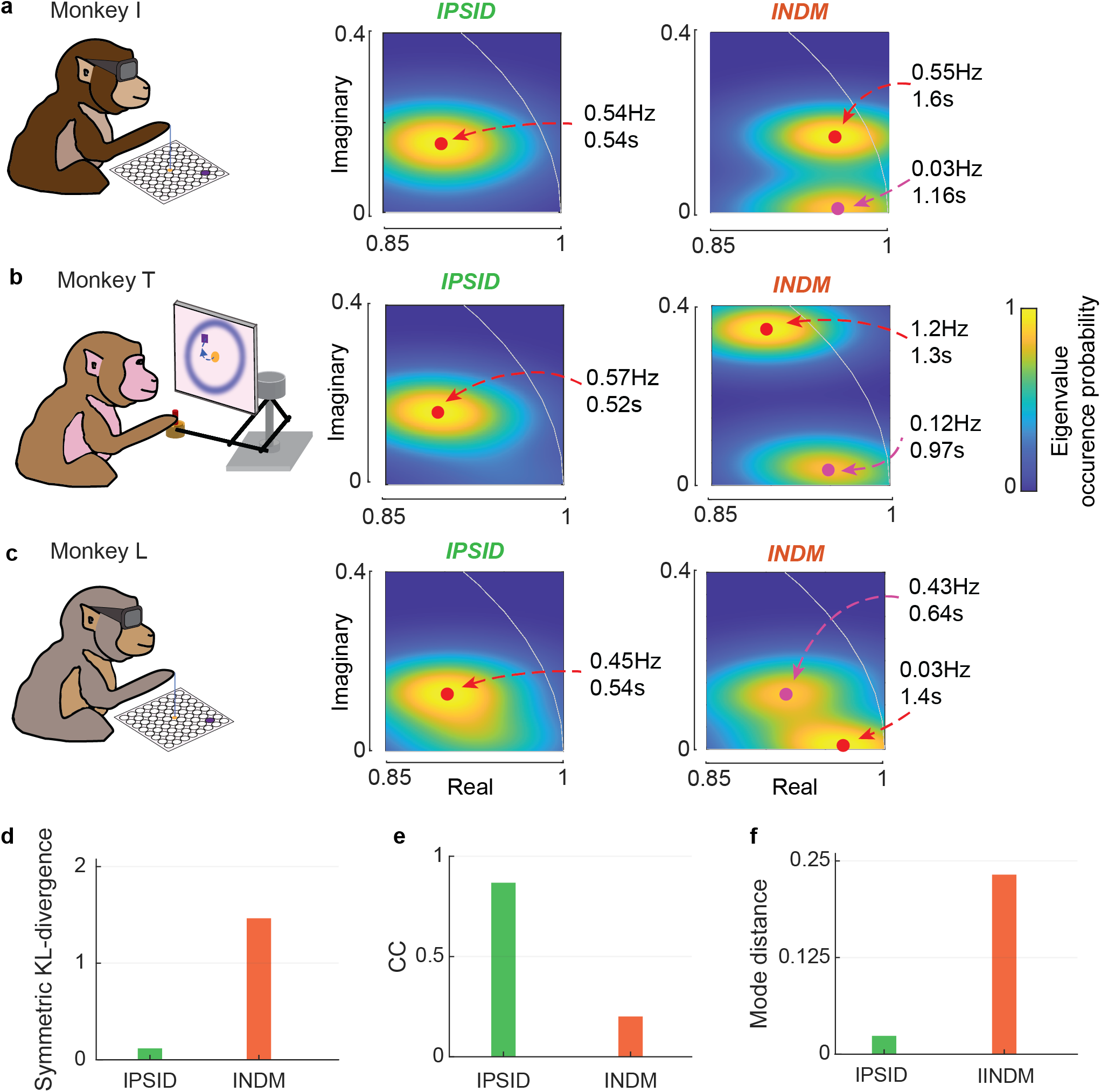
IPSID reveals largely similar intrinsic behaviorally relevant neural dynamics across three monkeys and two tasks from two independent datasets while INDM identifies different overall intrinsic neural dynamics. **(a)** Same as **Fig. 6b**, showing the eigenvalues learned for IPSID and INDM. **(b-c)** Similar to (a) for the second and third monkeys, respectively (taken from **Fig. 7** and **Fig. S11**). **(d)** Average pairwise symmetric KL-divergence between the eigenvalue probability mass functions of the three monkeys is computed for the IPSID/INDM results. **(e)** Average pairwise Pearson correlation coefficient (CC) of the probability mass functions of the three monkeys used to calculate the symmetric KL-divergence in (d). **(f)** Average pairwise distance between the mode (i.e., most probable eigenvalue location) of the probability mass functions of the three monkeys used to calculate the symmetric KL-divergence in (d). Lower KL-divergence/mode distance implies more similarity across monkeys, with a minimum possible value of 0. Higher CC implies more similarity across monkeys, with a maximum possible value of 1. Based on all three metrics, IPSID finds largely similar eigenvalues across tasks and animals whereas INDM finds eigenvalues that are different across tasks and animals.

We next found that models of neural dynamics learned by IPSID can generalize to other behavioral tasks that are not observed during training unlike methods that do not consider inputs. This is because by considering task instruction inputs, IPSID avoids learning models of neural dynamics that are confounded by the dynamics of instructions in a specific task. Indeed, models trained by IPSID on data from one task had minimal drop in behavior decoding performance when tested on data from a different task. In contrast, models learned by all other methods had significantly larger drops in behavior decoding performance in the other task (**Fig. S10**; *P* < 0.001; one-sided signed-rank; *n* = 10 simulations).

### Modeling task instructions as inputs reveals distinct intrinsic behaviorally relevant neural dynamics in non-human primate neural population activity

We next used IPSID to study intrinsic behaviorally relevant neural dynamics in two independent motor cortical datasets recorded from three monkeys (monkeys I and L from the first datasets and monkey T from the second dataset) during two distinct behavioral motor tasks with planar cursor movements (**Fig. 6a**, **Fig. 7a**). In the first dataset, which was made publicly available by the Sabes lab^45^, primary motor cortical (M1) population activity was recorded while two monkeys controlled a 2D cursor to reach random targets on a grid (**Fig. 6a**, **SI Methods**). The 3D position of the monkeys’ fingertip was tracked and the horizontal elements were passed on to control the cursor (**SI Methods**). In the second dataset, which was made publicly available by the Miller lab^46,47^, population activity from dorsal premotor cortex (PMd) was recorded while the monkey performed sequential reaches to random target positions on a plane (**Fig. 7a**, **SI Methods**). The cursor was controlled via a manipulandum that only allowed horizontal movements. For all subjects, we modeled the population spiking activity (**SI Methods**). We took the 2D position and velocity of the cursor as the behavior signal, and the timeseries of target positions as the input task instructions (**Figs. 6a, 7a**). We modeled the smoothed spike counts^3,13,39,48^ in all datasets as neural signals (**SI Methods**).

First, we found that IPSID revealed distinct intrinsic behaviorally relevant neural dynamics that were not found by other methods. Similar to our earlier simulation results (**Fig. 5**), this could be seen from the learned eigenvalues by IPSID that were different from those found by other methods (**Figs. 6b, 7b, S11b**). Second, eigenvalues found by PSID were far from those found by IPSID, whereas eigenvalues found by NDM were close to those found by INDM (**Figs. 6b**, **7b, S11b**). Note that IPSID/PSID focus on explaining the behaviorally relevant neural dynamics whereas INDM/NDM focus on explaining the overall neural dynamics regardless of relevance to behavior. Thus, the aforementioned result suggests that task instructions, which are taken as inputs in IPSID/INDM models, are highly informative of behaviorally relevant neural dynamics (seen from their effect on PSID vs IPSID), but are not very informative of the overall neural dynamics (seen from NDM and INDM results being similar). This is consistent with the vast body of work suggesting that neural dynamics relevant to any specific behavior may constitute a minority of the overall neural variance^5,6,19,32–39^.

In these analyses we used the additional steps in IPSID that were designed for scenarios in which some input dynamics may affect behavior through unrecorded regions/pathways (**Fig. S5**). However, we found that even without these additional steps, the average learned eigenvalues remained almost unchanged in one subject (**Fig. S12b**) and remained relatively similar in the other two subjects (**Fig. S12a,c**). This result could suggest, particularly in the former (**Fig. S12b**), that behaviorally relevant neural dynamics that were downstream of visual task instruction inputs were largely reflected in, or downstream of, the motor cortical recordings here. Overall, eigenvalues that were learned by IPSID were unique and were not learned by any of the other three methods. Having established their distinction, the next question was whether these distinct eigenvalues found by IPSID better describe the data, which we explored next.

### IPSID finds more accurate intrinsic behaviorally relevant neural dynamics in non-human primate neural population activity

We hypothesized that as in the simulation results (**Fig. 5**), the eigenvalues learned by IPSID are more accurate descriptions of the true intrinsic behaviorally relevant neural dynamics. We performed multiple evaluations to test this hypothesis. As a measure of closeness of two sets of dynamics, we computed the Kullback–Leibler (KL) divergence between the distribution of their associated eigenvalues (**SI Methods**).

First, we explored whether IPSID can mitigate the problem of learning eigenvalues that reflect input dynamics. We characterized the input dynamics by modeling the time series of task instructions as a linear state-space model (equation (3), **SI Methods**). From this model, we found that in all three subjects and in the two tasks, the eigenvalues representing the dynamics of input task instructions were close to those learned using NDM and PSID but not to those learned using IPSID (**Figs. 6b, 7b, S11b**). Indeed, in all subjects, the KL-divergence between the input dynamics and learned dynamics was much larger for IPSID compared with PSID which cannot consider inputs during learning (**Figs. 6c, 7c, S11c**). This result shows the success of IPSID’s novel algebraic operations in mitigating the influence of task instruction inputs on intrinsic dynamics unlike NDM and PSID.

Second, we demonstrated the success of preferential neural-behavioral modeling in the presence of input enabled by IPSID by comparing with INDM and NDM. In all three subjects, IPSID learned the intrinsic behaviorally relevant neural dynamics significantly more accurately than both INDM and NDM (**Figs. 6d**, **7d, S11d**). This was evident from comparing the cross-validated behavior decoding from neural activity for these methods (**Figs. 6d**, **7d, S11d**).

Third, we demonstrated the success of IPSID’s new algebraic operations in accounting for inputs in preferential neural-behavioral modeling by comparing it to PSID, which is preferential yet does not consider inputs. We found that by considering inputs, IPSID learns models that were significantly more predictive of neural dynamics compared to PSID in all three subjects, as evident by comparing the cross validated neural self-prediction accuracy across the two methods (**Figs. 6e, 7e, S11e**). These results held even if the feedthrough term *D_y_u_k_* in equation (2) – which reflects the effect of input on neural activity directly and not through the latent states *x_k_* – was discarded when predicting neural activity using IPSID (**Figs. 6e, 7e, S11e**). This analysis demonstrates that the better prediction in IPSID is due to its latent states being more predictive of neural dynamics rather than due to a static feedthrough effect of input on neural dynamics. Overall, these consistent results from three NHPs in two independent neural datasets with two different tasks suggest that IPSID can successfully dissociate intrinsic behaviorally relevant neural dynamics from other intrinsic neural dynamics and from measured input dynamics. Moreover, these results demonstrate that not considering task instruction sensory inputs when modeling neural activity can result in less accurate models of neural dynamics and confound conclusions about intrinsic dynamics, a problem that IPSID addresses (see also next section).

### IPSID uniquely revealed consistent intrinsic behaviorally relevant dynamics across three different subjects and two different tasks

While the specific task instructions are different in the two behavioral tasks in the independent datasets here – reaches to random targets on a grid vs. sequential reaches to random targets –, the two datasets also have similarities in terms of neural recordings and tasks. Specifically, both independent datasets have recordings from the motor cortical areas, and both involve cursor control tasks with targets on a 2D plane. We thus hypothesized that given these similarities, there may be similarities in the intrinsic behaviorally relevant neural dynamics across the two tasks and three subjects. To test this hypothesis, we compared the distribution of eigenvalues learned using IPSID across all pairs of the three subjects (**Fig. 8**) and quantified their average difference with three metrics: (1) symmetric KL divergence between eigenvalue distributions (**SI Methods**, **Fig. 8d**), (2) correlation coefficient (CC) between the probability mass functions of the eigenvalue distributions (**Fig. 8e**) (3) distance between the modes of the eigenvalue distributions, i.e., most probable locations (**Fig. 8f**). We also repeated these analyses for INDM.

We found that IPSID identified intrinsic behaviorally relevant dynamics that were strikingly similar across the two tasks and three subjects (**Fig. 8**). This result was clear from IPSID’s learned eigenvalues both qualitatively (**Fig. 8a-c**) and quantitatively based on the three abovementioned metrics (**Fig. 8d-f**). This similarity was despite the fact that the task instruction sensory inputs were distinct between the two tasks and that these recordings were from three different animals across two independent datasets. Also, even without its additional steps (**Fig. S5**, **Note S2**), IPSID still found largely similar eigenvalues across tasks and monkeys showing the robustness of this result, but the additional steps helped it reveal this similarity slightly more strongly (**Fig. S12d-f**).

We next studied the dynamics found by INDM. INDM aims to learn the overall intrinsic neural dynamics while IPSID aims to prioritize the learning of intrinsic behaviorally relevant neural dynamics. Interestingly, unlike IPSID, the dynamics found by INDM were much more distinct across the three monkeys (**Fig. 8**), as is clear both visually (**Fig. 8a-c**) and quantitatively (**Fig. 8d-f**). Moreover, as shown in the previous section, the more similar dynamics found by IPSID were also a more accurate description of intrinsic behaviorally relevant neural dynamics in each monkey (**Figs. 6d, 7d, S11d**). Together, these results suggest that while the overall intrinsic neural dynamics (as found by INDM) were different across these two planar motor tasks and three animals, the intrinsic behaviorally relevant neural dynamics were similar as revealed by IPSID. We propose that the similarity of the intrinsic behaviorally relevant neural dynamics may suggest that similar neural computations in the motor cortex underlie the planar cursor control tasks despite the differences between task instructions (i.e., inputs) and between animals.

IPSID was the only method that revealed the above finding about similar dynamics because it not only accounts for inputs (task instructions), but also prioritizes the learning of intrinsic behaviorally relevant dynamics over other neural dynamics in the presence of input – which is something INDM cannot do. Interestingly, this result is also consistent with our simulation study in **Fig. 5** in which IPSID was the only method that correctly found the fixed intrinsic behaviorally relevant dynamics regardless of task while other methods were confused by the task instructions and/or overall intrinsic dynamics. Thus, IPSID can help researchers explore and compare the intrinsic neural dynamics across different behavioral tasks without worrying about the specific structure of their task instruction inputs and how these inputs may confound their conclusions – e.g., mitigate the confound that similarity or lack thereof in dynamics may simply be due to input comparisons across tasks.

Together, these results highlight that the new algebraic operations in IPSID can lead to both more accurate models and new useful scientific insight. These results also demonstrate that even though a comprehensive measurement of all inputs to a given brain region is typically experimentally infeasible, even incorporating partial input measurements (task instruction sensory inputs in this case) can already yield new insights into neural computations across different tasks and subjects.

## Discussion

We developed IPSID, a novel method that provides the new capability to perform preferential dynamical modeling of neural-behavioral data in the presence of measured inputs. In the IPSID formulation, a dynamical model of neural activity is learned by accounting for measured input, neural, and behavioral data simultaneously, and the learning of intrinsic behaviorally relevant neural dynamics is prioritized over other intrinsic dynamics. By doing so, IPSID can dissociate intrinsic behaviorally relevant dynamics not only from other intrinsic dynamics, but also from the dynamics of measured inputs such as task instructions or recorded activity of upstream regions. We demonstrated that without IPSID, dynamics in measured inputs to a given brain region or other intrinsic neural dynamics may be incorrectly identified as intrinsic behaviorally relevant neural dynamics within that brain region and thus confound conclusions. Indeed, in the neural data from monkeys, we showed that task instructions can act as such confounding inputs. IPSID can analytically account for such measured inputs to reveal distinct and more accurate intrinsic behaviorally relevant neural dynamics compared with existing approaches even when they considered input (as in INDM). By doing so, IPSID also provided useful scientific insights about intrinsic neural dynamics of behavior across different tasks and animals, which were not found by other methods.

IPSID could allow future studies to more easily design and compare across tasks without worrying about the temporal structure of task instruction inputs and how their reflection in neural activity may be misinterpreted as intrinsic neural dynamics. We showed this potential with experiments where a simulated brain with fixed intrinsic dynamics performed different cursor control tasks. We showed that sensory inputs in the form of task instructions could lead to learning intrinsic dynamics that incorrectly appeared task-dependent and different across tasks. IPSID addressed this issue and was the only approach that correctly found the intrinsic behaviorally relevant neural dynamics regardless of the task. Consistently, in the real motor cortical datasets and by modeling the task instructions as sensory inputs, IPSID not only learned the intrinsic behaviorally relevant neural dynamics more accurately, but also was the only method that revealed their similarity across tasks and animals.

Unexpectedly, despite differences in animals and in motor tasks and their instructions across the motor cortical datasets, we found similar intrinsic behaviorally relevant dynamics in all three animals across both tasks/datasets using IPSID. In contrast, INDM found that the dominant overall intrinsic dynamics were different across tasks and animals. This result may suggest that motor cortical regions across different animals could have different intrinsic dynamics overall, but the part of their intrinsic dynamics that is engaged in arm movements to control 2D planar cursors may have similarity. These similar dynamics may suggest that similar intrinsic neural computations in the motor cortex underlie the performance of these two different planar cursor-reaching tasks. Prior work has found similarities in static projections of neural activity^49,50^ across subjects^50^ or tasks^49^, but these prior works have not modeled temporal dynamics (e.g., eigenvalues) and have not disentangled the effect of task instruction input dynamics on the observed similarity. Thus, IPSID provides a new useful tool to explore whether such observed similarities reflect input dynamics or are intrinsic.

When the activity of some upstream brain regions that have inputs to the recorded region^27,31,51–53^ is not measured, the learned intrinsic dynamics could also partly originate from these other regions. In the motor cortical datasets here for example, neural dynamics in upstream regions such as visual cortex—which is involved in processing the sensory input and passing it to other regions along the visual-motor pathway—may also be reflected in the learned intrinsic motor cortical dynamics. Taking the sensory instructions as input can, to some extent, account for the dynamics of inputs from these upstream visual areas. Similarly, a sensory input that is not measured or accounted for, for example the sunrise-sunset cycles during chronic recordings, may confound the modeled neural dynamics of a specific behavioral or mental state such as mood (e.g., in the form of circadian rhythms)^54,55^. Thus, recording activity from more upstream regions and measuring more sensory inputs can allow IPSID to analytically consider more comprehensive inputs during modeling to better discover intrinsic behaviorally relevant dynamics. As it is mostly experimentally infeasible to identify and record all inputs to a given brain region, a complete disentanglement of intrinsic dynamics from all input dynamics to a region becomes impractical. This experimental limitation is thus a fundamental limit on methodological efforts aimed at disentanglement. Thus, one still needs to interpret the results cautiously by noting that only dynamics of measured inputs are being disentangled from intrinsic dynamics. Nevertheless, our results show that even this partial disentanglement can lead to more accurate models and to new useful insights compared to alternative models which either do not consider measured inputs, or consider measured input but not behavior during learning.

Here, we address the challenge of preferential modeling of neural-behavioral data with measured inputs, which has been unresolved. For non-preferential modeling of neural data on its own and when inputs are not measured, prior studies have looked at the distinct problem of separating the recorded neural dynamics into intrinsic dynamics and a dynamic input that is inferred^12,56,57^. This decomposition is typically done by making certain a-priori assumptions about the input such that inputs can be inferred, for example that input is constrained to be considerably less dynamic than intrinsic neural dynamics, or that input is sparse or spatiotemporally independent^12,56^. In addition to preferential neural-behavioral modeling with measured inputs, which is addressed here, future work can extend preferential modeling to also incorporate similar input inference approaches, which could be complementary to IPSID. For example, such input inference approaches can help further interpret the intrinsic behaviorally relevant dynamics extracted by IPSID and hypothesize which parts of them could be due to unmeasured inputs. The results from such input inference efforts can depend on the a-priori assumptions made regarding the input, since mathematically both extremes are plausible when inputs are not measured: all neural dynamics could be due to input from another area or they could all be intrinsic. For this reason, validating the inferred inputs from these inference approaches against actually measured inputs is an important step^12,53,56,57^. Such validation is also important because the underlying dynamics and inputs can have potential nonlinearities, thus making the inference of unmeasured inputs challenging or infeasible due to the potential unidentifiability in nonlinear systems^58^.

One main contribution here is to formulate and highlight the problem of how intrinsic neural dynamics underlying a specific behavior can be confounded by both input dynamics and other intrinsic neural dynamics. We formulated this disentanglement problem that simultaneously involves measured input, neural, and behavioral data during learning, and derived IPSID as a new analytical solution based on subspace identification. By comparing with INDM and a block-structured numerical optimization approach (**Figs. 3–4**), we showed that two new capabilities in IPSID are critical for disentanglement: prioritized learning of intrinsic behaviorally relevant dynamics via the new two-stage learning operations with inputs, and dissociating those behavior dynamics that are due to input but not reflected in the neural recordings from those that are via the additional analytical steps (**Fig. S1**, **Fig. S5**). Prior works have proposed enforcing block-structure on linear dynamic models and developed Expectation-Maximization algorithms for fitting them^59,60^. But these studies have distinct goals and thus do not address the input disentanglement problem, or the behaviorally relevant dissociation problem addressed here. As such, they also do not enable the above two new capabilities enabled by IPSID that are critical for solving these problems. Future work can utilize the ideas developed here for enabling the IPSID capabilities in order to develop alternative numerical optimization solutions to the formulated disentanglement problem.

In addition to sensory inputs or activity in other brain regions, the input could also be any external electrical or optogenetic brain stimulation, for example in a brain-machine interface (BMI). Developing novel closed-loop stimulation treatments for mental disorders such as depression^61,62^ hinges on building dynamic models of neural activity that satisfy two criteria: (i) describe how mental states are encoded in neural activity^61,62^; (ii) describe the effect of electrical stimulation on the neural activity^28,62,63^. The approach developed here enables learning of models that satisfy both criteria. First, by prioritizing behaviorally relevant dynamics, models accurately learn the neural dynamics relevant to behavioral measurements of mental states (e.g. mood reports in depression^61^). Further, this prioritization enables the learned models to have lower-dimensional latent states, which is important in developing robust controllers^64^. Second, the models can explicitly learn the effect of external electrical stimulation parameters on neural activity^28,63^.

Here we used continuous valued variables with Gaussian distributions to model neural activity, as has been done extensively in prior works modeling local field potentials (LFP)^14,19,30,44,61,65,66^ and spike counts^7,19,67,68^. However, recent works suggest that modeling spike counts as Poisson distributed variables^8,12,69–72^ can improve BMI performance^70,71^. Thus, an interesting direction is to extend the method to support Poisson distributed neural observations, or support simultaneous Gaussian and Poisson neural observations for multiscale modeling of neural modalities such as LFP and spikes together^16,44,65,73–75^. Supporting general nonlinearities in the intrinsic dynamics and their relation to behavior is another interesting future direction^20,41^. Finally, developing adaptive extensions that update the dynamical latent state model to adapt to non-stationarities in neural signals or to stimulation-induced plasticity^43,76–79^ will be important for BMIs and for studying learning and plasticity and their effect on intrinsic behaviorally relevant dynamics.

In conclusion, we provide a new analytical method for preferential dynamical modeling of neural-behavioral data that can account for measured inputs—whether sensory input, neural input from other regions, or external stimulation. We show the importance of doing so for correct interpretation of neural computations and dynamics that underlie behavior, for accurate modeling of intrinsic neural dynamics, and for gaining useful insights about neural computations of behavior across different tasks and subjects. These results and the new preferential modeling approach have important implications for future neuroscientific and neuroengineering studies.

## Acknowledgements

This work was partly supported by NIH R01MH123770 and DP2MH126378 (M.M.S) and USC Annenberg Fellowship (O.G.S). We sincerely thank the Sabes lab at the University of California San Francisco^45^ and the Miller lab at Northwestern University^46,47^ for making the NHP datasets^45–47^ that we used here publicly available.

## Author contributions

P.V., O.G.S., and M.M.S developed the algorithms, analyzed data, and wrote the manuscript. M.M.S. supervised the study.

## Supplementary Information for

### Supplementary Information Methods (SI Methods)

#### Non-preferential neural dynamic modeling with and without input

Neural population activity exhibits rich temporal structures^1–22,26,23,25,24^. Neural dynamical modeling aims to describe all such temporal structures in neural activity^2–5,7,8,10–14,16,21,23,24,26,61,65^ without prioritizing the learning of dynamics related to any particular behavior, which is why we also refer to it as non-preferential modeling. In this section we provide a brief overview of linear neural dynamical modeling with and without consideration of inputs, referred to respectively as INDM and NDM^2,7,8,14,16,19,61,65^. In NDM, neural population activity is modeled in terms of a latent state as

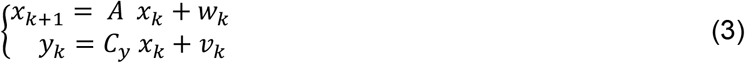

where as before, 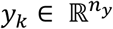 and 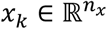 represent the neural activity and the latent state of the neural population, respectively. 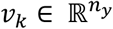 represents noises in neural activity and 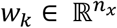 represents excitations that drive the latent state, with their covariances defined as in equation (5) below. Given that in the NDM formulation inputs are not explicitly modeled, the excitations of the latent state represented by *w_k_* will capture a mixture of both the measured inputs and all other excitations that originate within the recorded brain region or from other brain regions. However, since *w_k_* is modeled as noise and not a dynamical system itself, the NDM model has to capture any temporal structure in inputs via the dynamics of the latent state *x_k_*, which is quantified through the state transition matrix *A*. Thus, when inputs are temporally structured, the state dynamics—which are subsequently reflected in the neural activity *y_k_*—may incorrectly also incorporate the structured dynamics that exist in inputs, such as sensory inputs in the form of movement targets visually presented on a screen^1,9,12,27,30^ (we also show this in our results). As overviewed next, when inputs are measured, although not commonly done, INDM methods can be used to incorporate them in non-preferential modeling.

In INDM, measured inputs are incorporated into equation (3) as

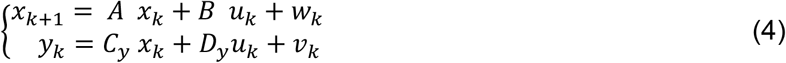

where 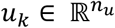 is the measured input to the neural population, e.g. inputs from other brain regions or sensory inputs. Unlike equation (3), in equation (4), the term *u_k_* explicitly represents measured inputs to the recorded brain region and thus the state dynamics reflected in *A* no longer need to describe the dynamics of *u_k_*; rather, state dynamics solely need to describe the intrinsic dynamics of the brain region in response to measured (*u_k_*) and unmeasured excitations (*w_k_*). Thus, this approach can dissociate the input dynamics form the intrinsic dynamics. While not commonly used in neuroscience, INDM has great utility in system identification^40,80^. However, INDM does not allow for preferential modeling of shared dynamics between two signals, such as neural activity and behavior. Here we address the unresolved challenge of modeling the effect of input within preferential modeling of two signals together (e.g., neural population activity and behavior).

#### Enabling the modeling of input effects in preferential dynamical modeling

##### Model formulation

We develop a new method termed IPSID to enable dissociating the effect of input in preferential dynamical modeling of neural-behavioral data, which has not been possible to date. Unlike non-preferential dynamical modeling—e.g., NDM/INDM and other approaches^2–4,7,8,10–14,16^—, which models the dynamics of a single signal such as neural population activity, preferential modeling dissociates the shared dynamics between two signals, such as neural population activity and behavior, and prioritizes the learning of these shared dynamics^19^.

So far, input effects cannot be considered in preferential dynamical modeling, even when input is fully measured. Here we develop a preferential dynamical method that can achieve this goal and dissociate input dynamics and intrinsic dynamics. Specifically, the method simultaneously achieves two goals: It dissociates the intrinsic dynamics that originate in the recorded region from input dynamics that are simply reflected in the recorded regions but do not originate there (e.g., input from upstream regions or sensory feedback). Second, it dissociates and prioritizes the learning of intrinsic neural dynamics that are relevant to the behavior of interest from other intrinsic neural dynamics.

This new method jointly models neural activity 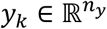, behavior 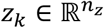, and the effect of input 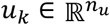 on them with the general linear state-space formulation in equation (1). In equation (1), 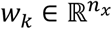 and 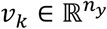 are taken to be zero-mean white noises that are independent of *x_k_*, i.e. 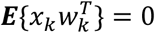 and 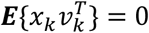 with the following cross-correlations:

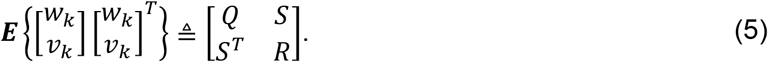

The latent state *x_k_* describes all intrinsic neural dynamics including those related to the given behavior and those unrelated to it. It can be shown^19,80^ that equation (1) can always be written in an equivalent basis as

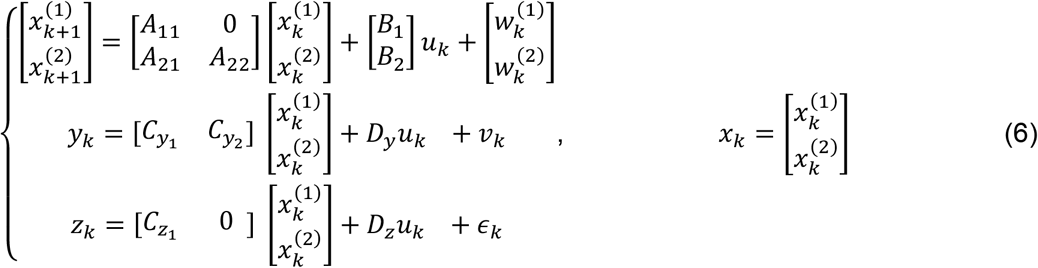

where 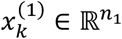 and 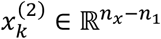 denote the behaviorally relevant and the other dimensions of *x_k_*, respectively. In IPSID, we directly learn the model in the basis shown in equation (6) in a way that prioritizes behaviorally relevant states. *n*_1_ is the smallest latent state dimension that is sufficient for explaining all the dynamics of behavior *z_k_* that are encoded in neural activity *y_k_*. The parameter set (*A*, *B*, *C_y_*, *C_z_*, *D_y_*, *D_z_*, *Q*, *R*, *S*) fully specifies the model in equation (1) and equivalently in equation (6), except for dynamics of *ϵ_k_*, which represent behavior dynamics that are not encoded in neural activity. As explained in **Note S2**, when desired (e.g., see **Fig. S8a**), dynamics of *ϵ_k_* can be modeled separately by modeling the residual behavior after equation (1) is used to predict behavior^19^ (**Fig. S5c**).

##### Learning model parameters using IPSID

In the learning problem, given neural, behavior, and input time-series – denoted by {*y_k_*: 0 ≤ *k* < *N*}, {*z_k_*: 0 ≤ *k* < *N*}, and {*u_k_*: 0 ≤ *k* < *N*}, respectively –, and given the desired dimension of the latent state *n_x_*, and the desired number of behaviorally relevant latent states *n*_1_, the aim is to learn all model parameters in equation (6) while prioritizing learning of behaviorally relevant states. To do this, during training, we first extract the intrinsic latent states directly using the neural activity, behavior and input training data; we then identify the model parameters using the extracted intrinsic latent states. The details are provided in **Note S1**. Here, we briefly explain the algorithm and the intuition behind it.

IPSID extracts the latent states from training data in two stages (**Fig. S1**). In stage 1, the subspace spanned by the behaviorally relevant latent states 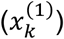 is extracted with *priority* by projecting future behavior (*Z_f_*) onto past neural activity (*Y_p_*). However, when there exist measured inputs, the input dynamics also affect future behavior. Thus, an orthogonal projection of future behavior onto past neural activity (as is used in our prior work on PSID^19,41^) would give a mixture of the subspaces spanned by the intrinsic behaviorally relevant latent states and the input. This means that the learned states will not just reflect the intrinsic behaviorally relevant neural dynamics as these intrinsic dynamics cannot be dissociated from input dynamics. To extract the subspace associated with intrinsic behaviorally relevant latent states, we have to devise distinct algebraic operations in IPSID.

We thus devise these algebraic operations to perform an oblique (i.e., non-orthogonal) projection of future behavior onto past neural activity and past input along the subspace spanned by the future input (**Fig. S1**, **Fig. S6c**). Note that the use of oblique projections in IPSID instead of orthogonal projections is not as simple as replacing one operation with another; rather, this key change has broad consequences throughout the rest of the model learning operations that are appropriately accounted for in IPSID. For example, the learning of input parameter *B* (see equation (1)) has no equivalence in prior works that do not consider input (e.g., PSID) and requires distinct operations. The oblique projection ensures that the result of the projection is orthogonal to the subspace spanned by the future input, thus excluding behavior dynamics that can be directly attributed to future input dynamics rather than intrinsic neural dynamics (**Note S1**).

In an optional stage 2, we devise additional algebraic operations to extract the subspace spanned by any remaining latent states 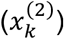 via an oblique projection from the residual future neural activity—the part unexplained by the extracted behaviorally relevant states 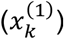—onto past neural activity and past input, again along the subspace spanned by the future input (**Fig. S1**). Finally, given the fully extracted subspace of the latent states (either only from stage 1 or by concatenating the results from both stages), we learn all model parameters based on equation (1) via least squares as detailed in **Note S1**.

##### Special cases of IPSID

IPSID addresses two challenges simultaneously by allowing for input incorporation in preferential dynamical modeling of neural-behavioral data together. First, it dissociates intrinsic and measured input dynamics. Second, it dissociates intrinsic behaviorally relevant dynamics from other intrinsic neural dynamics and prioritizes the learning of the former. Special cases of IPSID cover prior methods developed by us or others, which only address one of the challenges that IPSID addresses. Briefly, when not modeling input (equivalently when assuming *B*, *D_y_*, *D_z_* are zero in equation (1)), IPSID reduces to PSID in our prior work^19^. In this case, input dynamics may be learned as part of the intrinsic dynamics in the model and thus the first challenge of dissociating the dynamics of measured input and intrinsic dynamics will not be addressed. Alternatively, when behavior is not considered during modeling, i.e., when *n*_1_ = 0 such that all latent states are extracted using the second stage of IPSID, IPSID reduces to prior neural dynamical modeling with input (i.e., INDM)^40,80^, which is formulated by equation (4). In this case, intrinsic behaviorally relevant dynamics are not dissociated or prioritized compared with other intrinsic neural dynamics, and thus the second challenge will not be addressed. Finally, if inputs are not considered and only the second stage is used, IPSID reduces to prior neural dynamical modeling (NDM) formulated by equation (3), which does not address either of the two challenges addressed by IPSID.

##### Learning using numerical optimization with block-structured parameters

To compare with IPSID and show the benefits of its two-stage learning method in prioritized learning of intrinsic behaviorally relevant dynamics, we also implement an alternative approach for fitting the same model using standard numerical optimization. In this approach we use numerical optimization^81^ to learn all model parameters by maximizing the neural-behavioral data log-likelihood while imposing the same block-structure as is defined in equation (6). To do so, we use the recurrent neural network class^†^ in TensorFlow^82^ v2.5 to implement a linear recurrent neural network with the computation graph corresponding to equation (6). This approach uses gradient descent via error backpropagation to fit the parameters of the recurrent neural network. Parameters are learned while maximizing the log likelihood of full output data, i.e., both neural activity and behavior [*y_k_*, *z_k_*]. We use minibatch size of 32 and run the numerical optimization on the same CPUs as was done for IPSID to enable a fair comparison in terms of the training time (**Fig. S9**). While the sequential nature of recurrent neural networks limits parallelization options during training^83^, it is possible that other implementations (e.g., using a framework other than TensorFlow^82^) or computational tricks^84^ may have different results and may lead to faster learning. Note that while this numerical optimization enforces the same block-structure, it cannot prioritize the learning of intrinsic behaviorally relevant dynamics and does not dissociate behavior dynamics not reflected in the neural recordings. As we show in results, both these new capabilities are critical for disentanglement.

##### Additional steps to add IPSID support for scenarios where neural recordings do not cover/reflect all downstream regions of the input

We also add additional optional steps to IPSID to support scenarios where some downstream regions of input that affect behavior are not covered by the recorded neural activity. In these scenarios, which we formulate as in equation (2), input may affect behavior through paths that are not reflected in the recorded neural activity. We thus add a new type of latent state in equation (2), denoted by 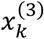, to describe such paths as latent states that are affected by input and can affect behavior but do not contribute to generation of neural activity, i.e., columns corresponding to these states are zero in *C_y_* and the noises driving them (i.e., 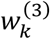) are uncorrelated with the noises that drive the other latent states (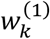 and 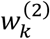). To dissociate these states from other states in IPSID, we perform two additional steps (**Note S2**, **Fig. S5**).

First, before performing the two-stage IPSID learning process described earlier (**Note S1**), we use the IPSID second stage alone to build a model for all intrinsic neural dynamics in terms of a high-dimensional latent state, denoted by 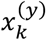 in **Note S2** and **Fig. S5**. We next project the behavior onto the extracted latent state 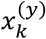 (with the result denoted by 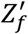 in **Note S2** and **Fig. S5**); doing this projection removes any elements of behavior that are not encoded in neural activity (**Fig. S5a**). We then proceed with IPSID stages 1 and 2 as before but with the projected behavior signal (i.e., 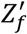) used in the modeling rather than the original behavior signal (**Fig. S5b, Fig. S6a**); using this projected behavior signal ensures that behavior dynamics that are not encoded in the neural recordings are not included in the first set of states 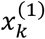.

Second, if learning of behavior dynamics that are predictable from input but are not reflected in neural activity is of interest, we perform an additional step to learn a model for an additional latent state 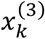 to describe such dynamics. To do this, we first subtract from behavior its prediction from the past neural activity and past input using the already learned model that consists of 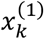 and 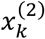. This gives us a residual behavior signal. We then apply the IPSID second stage alone to this residual behavior to model its dynamics and the effect of input on it in terms of a latent state 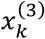, with a formulation akin to equation (4) but with *z_k_* as output of the second line. This gives 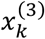 because 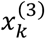 summarizes the direct effect of input on behavior dynamics that are not reflected in the recorded neural activity. We then put the model learned for 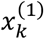 and 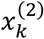 together with the model learned for 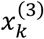 to get a full model in the form of equation (2). Note that the model for 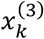 has no neural observation. Thus, in the final overall model, the columns of *C_y_* corresponding to 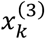 will be zero. Consequently, neural activity *y_k_* makes no contribution to the estimation of 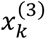 (see equation (7) below) in the overall model, resulting in the estimated 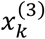 being forward predicted purely using input *u_k_*. This concludes the learning of the full model as formulated in equation (2). The details are provided in **Note S2**.

##### Extracting latent states and predicting neural activity and behavior using the learned model

Given the model in equation (1), the prediction of behavior and neural activity given past neural activity and inputs are obtained using the well-known recursive Kalman filter^40,80^. Thus, once the model is learned, in test data, we extract the latent states by applying the Kalman filter associated with the identified model parameters to the neural activity and input as

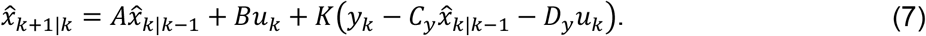

Here 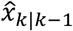 denotes the latent state estimated at time step *k* using neural activity and input up to time step *k* – 1 (*y_n_* and *u_n_* for 0 ≤ *n* < *k*), with 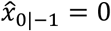 taken as the initial state. *K* denotes the steady state Kalman gain^40,80^, which can be computed as *K* = (*APC^T^* + *S*)(*CPC^T^* + *R*)^−1^ where *P* is the solution to the following steady-state Riccati equation: *P* = *APA^T^* + *Q* – (*APC^T^* + *S*)(*CPC^T^* + *R*)^−1^(*APC^T^* + *S*)^*T*^. Note that behavior is not looked at when extracting the latent states using the Kalman filter (it is only looked at during model training in the training set). Given the estimated latent states 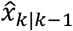, we next compute the self-prediction of neural activity 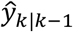 and the decoding of behavior 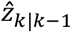, both given the past neural activity (*y_n_* for 0 ≤ *n* < *k*) and the past and current inputs (*u_n_* for 0 ≤ *n* ≤ *k*) as

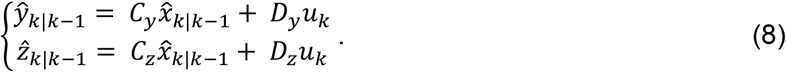

#### Performance measures: behavior decoding and neural self-prediction

To evaluate the leaned models and as a measure of how well the behaviorally relevant neural dynamics are learned, we compute the accuracy of decoding behavior (equation (8)). We perform modeling within a 5-fold cross-validation. As the performance measure, we compute the correlation coefficient (CC) between the decoded and true behavior time-series in the test data, averaged across the data dimensions, as the performance measure. Similarly, to evaluate how well the neural dynamics are learned in general, irrespective of their relevance to behavior, we compute the accuracy (in terms of CC) of predicting neural activity one-step-ahead using past neural activity and past and current inputs (equation (8)), which we refer to as neural self-prediction.

#### General simulations with random models

To validate IPSID with numerical simulations, we generate random models and confirm that IPSID can correctly learn these models when provided with training data. We generate the random models as follows. First, we select the dimensions of *y_k_*, *z_k_* and *u_k_* with uniform probability from ranges 5 ≤ *n_y_*, *n_z_* ≤ 10 and 1 ≤ *n_u_* ≤ 4. We select the full latent state dimension *n_x_* uniformly from 1 ≤ *n_x_* ≤ 10 and then we select the dimension of the state that derives behavior, i.e., *n*_1_, uniformly from 1 ≤ *n*_1_ ≤ *n_x_*. We generate the state transition matrix *A* based on its eigenvalues, which we randomly generate in complex conjugate pairs drawn with uniform probability across the unit disk. We randomly select a subset of *n*_1_ complex-conjugate eigenvalues as the behaviorally relevant eigenvalues to be placed in the top-left *n*_1_ × *n*_1_ submatrix of *A* (i.e., *A*_11_ in equation (6)). We generate matrices *B*, *C_y_*, and *C_z_* randomly with standard normal distribution for each of their elements, and then we set the right block of *C_z_* (after the first *n*_1_ columns) to zero as in equation (6)). For our simulation analyses in **Fig. 2a-b**, **Figs. S2–S4**, both *D_y_* and *D_z_* are randomly generated non-zero matrices. In all other figures, we take *D_z_ =* 0, and in **Figs. 1,3,5,6**, we additionally take *D_y_* = 0. Finally, we generate a random positive-semi-definite square matrix with *n_x_* + *n_y_* rows as the noise covariances *Q*, *R*, and *S* per equation (5). We then select two random numbers between 0.1 to 10 (with uniform probability in log scale) and scale the state and observation noises with these numbers to provide a wide range of relative state and observation noise values; we reflect that scaling in covariances *Q*, *R*, and *S*.

With a similar approach, we generate two other random models but this time without input: one as the behavior noise model to generate the behavior dynamics not present in neural activity, i.e. *ϵ_k_*; and, one as the input model to generate the input *u_k_*. The output dimension in these models is selected consistent with the behavior and input dimensions in the main model, respectively. We select the dimension of the latent state in the behavior noise model uniformly in 1 ≤ *n_x_ϵ__* ≤ 10 and that of the input model uniformly from 1 ≤ *n_x_u__* ≤ 4. Finally, to cover a diverse range of signal to noise ratios (i.e., signal *C_z_x_k_* over noise *ϵ_k_*) for behavior, we select a random number between 1 and 100 (with uniform probability in log scale) and scale rows of *C_z_* such that the ratio of the signal s.d. to noise s.d. for each behavior dimension becomes the selected random number. Note that the eigenvalues of the state transition matrix *A_u_* in the input model are representative of the input dynamics and may incorrectly be learned as intrinsic dynamics by methods that do not consider input (NDM/PSID) (**Fig. 2** and **Fig. S3**). To simulate scenarios in which input affects behavior through pathways that are not reflected in the recorded neural activity (**Fig. 3, Figs. S7–S8**), we add an input term *B′u_k_* to the state equation of the model that generates *ϵ_k_* with a *B*′ parameter that is non-zero only in a subset of rows; this way, the input affects a random subset of dimensions of the states that generate *ϵ_k_*, effectively changing their role to that of 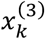 in equation (2).

Given the model parameters, a time series realization with *N* data points can be generated with the following procedure. First, the input time series *u_k_* is generated by drawing *N* Gaussian noise samples with the covariances given in the input model (similar to *Q*, *R*, and *S* in equation (5)) and iterating through the state-space equation (similar to equation (3), but with *y_k_* renamed to *u_k_*). This gives the *N*-sample input time-series {*u_k_*: 0 ≤ *k* < *N*}. Similarly, a realization from the behavior noise model is generated as the *N*-sample time-series {*ϵ_k_*: 0 ≤ *k* < *N*}. Next, *N* Gaussian noise samples are randomly generated for noise covariances in the main model (*Q*, *R*, and *S* in equation (5)). These noise samples along with *u_k_* and *ϵ_k_* are used to iterate through equation (1) and produce the behavior and neural activity time series { *y_k_,z_k_*: 0 ≤ *k* < *N*}. In all cases, the initial state in the state-space model iterations is taken to be *x*_−1_ = 0. Given the generated neural activity, behavior and input time series, we fit a model using (I)PSID and (I)NDM algorithms with horizon parameter of *i* = 5 (**Note S1**).

#### Performance measures for learning of model parameters and eigenvalues in numerical simulations

The model in equation (1) can be written with infinitely many different but equivalent sets of parameters that all give rise to the exact same statistics for neural and behavior observations *y_k_* and *z_k_*. Thus, to evaluate the parameter learning performance, we need to consider all equivalent sets of parameters for the identified model. An equivalent model to a given model can be obtained by a change of the latent state basis (also known as a similarity transform), which can be obtained by multiplying the latent state with an invertible matrix. Thus, to compare the identified and true models, we first solve an optimization problem to find the similarity transform that makes the basis of the identified model as similar as possible to the true model, and then compute the difference between the identified and true model parameters. Just to find this similarity transform, we generate a new realization with *q* = 1000*n_x_* samples from the true model, and then extract latent state 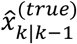 and 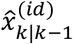 using the steady-state Kalman filter associated with the true and identified models (equation (7)), respectively. We then find the optimal similarity transform that minimizes the mean squared error between the two sets of latent states as

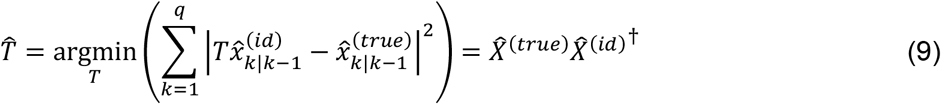

where 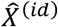 and 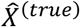 are *n_x_* × *q* matrices whose *k*th columns are constructed of 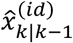 and 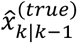 respectively. We apply the similarity transform^80^ associated with 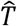 (i.e., the transform that changes the states from 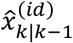 to 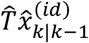) to the identified model parameters and then quantify the identification error of each parameter as

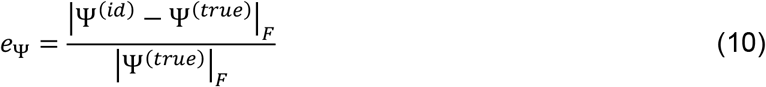

where |.|_*F*_ denotes the Frobenius norm of a matrix Ψ, which for any matrix Ψ = [*ψ_ij_*]_*n*×*m*_ is defined as

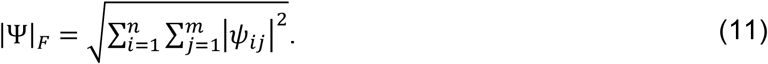

In addition to computing the identification error for all model parameters, we also compute the error in how accurately the eigenvalues of the state transition matrix *A* are learned. This metric is an additional indication of how well the dynamics are learned since the eigenvalues of *A* affect the transfer function from the input to the states and to the neural activity (**Fig. 1b**), and determine the frequency and decay rate with which the latent state responds to excitations from the state noise *w_k_* or from inputs *u_k_* (equation (1))^85^. To measure how well the intrinsic behaviorally relevant dynamics are learned, we evaluate how well the eigenvalues of the behaviorally relevant block of *A* (i.e., *A*_11_ in equation (6)), referred to as behaviorally relevant eigenvalues, are learned. These behaviorally relevant eigenvalues are obtained for each method as follows.

IPSID/PSID learn the model in the form of equation (6). So for these IPSID/PSID models, we simply compute the eigenvalues of *A*_11_ (which has dimension *n*_1_) and find their minimum normalized distance from the true behaviorally relevant eigenvalues by placing the eigenvalues in a vector and computing the error per equation (10), e.g., in **Fig. 2**. Note, in simulations, all methods know the dimension of the true behaviorally relevant latent states denoted by *n*_1_ and the dimension of the latent state in the input model (i.e., dimension of input dynamics) denoted by *n_x_u__* (see above). For PSID and because it does not consider the input and thus cannot dissociate intrinsic and input dynamics, in its first stage we use a state dimension equal to *n*_1_ + *n_x_u__*, so that its first stage has enough state dimensions to capture both the input dynamics and the intrinsic behaviorally relevant neural dynamics. Then we take the top *n*_1_ state dimensions that are best for decoding behavior as the behaviorally relevant states and evaluate the distance of their associated eigenvalues to the true behaviorally relevant eigenvalues (we refer to this procedure as model reduction, see also below for INDM/NDM).

INDM/NDM do not dissociate the behaviorally relevant latent states. Thus, for INDM/NDM, to find these latent states and their associated eigenvalues in the learned models, we proceed as follows: we first perform a similarity transform (using MATLAB’s bdschur command followed by the cdf2rdf command) to find an equivalent model with block-diagonal *A*. We next use the Kalman estimated latent states in each block to predict behavior. We then sort all blocks in descending order of their behavior decoding performance. Finally, we take the top *n*_1_ eigenvalues associated with the blocks with the best decoding performance as the behaviorally relevant eigenvalues in the identified model. We next compute the error in these eigenvalues using equation (10) as was explained earlier for IPSID/PSID (**Fig. 2**). We refer to this procedure as model reduction; for example, we can fit a high-dimensional INDM model and then reduce its dimension by finding those dimensions/eigenvalues that are best in decoding behavior (e.g., see **Fig. 2c**).

#### Motor task simulations

We devise a numerical simulation of a brain performing various cursor control tasks (**Fig. 5**). We use this simulation to demonstrate the role of sensory task instructions as inputs to the brain that affect neural dynamics and can confound the learned models of intrinsic neural dynamics. We modeled the brain with two components (**Fig. 5a**): (i) A linear state-space model (LSSM) in the form of equation (12) with 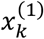 corresponding to the 2D position and velocity of the cursor (overall a 4D state; 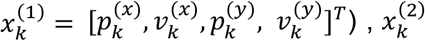 a 2D latent state corresponding to neural dynamics unrelated to behavior, and 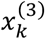 a 2D latent state corresponding to additional input driven dynamics in behavior that are absent in neural activity. (ii) An optimal feedback controller (OFC) that tries to control the part of the latent state in the LSSM that represents the cursor kinematics (i.e., 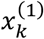) such that the cursor moves to targets presented via the task instructions^42–44,70^. The latent brain state *x_k_*, neural activity *y_k_* and behavior *z_k_* evolve as

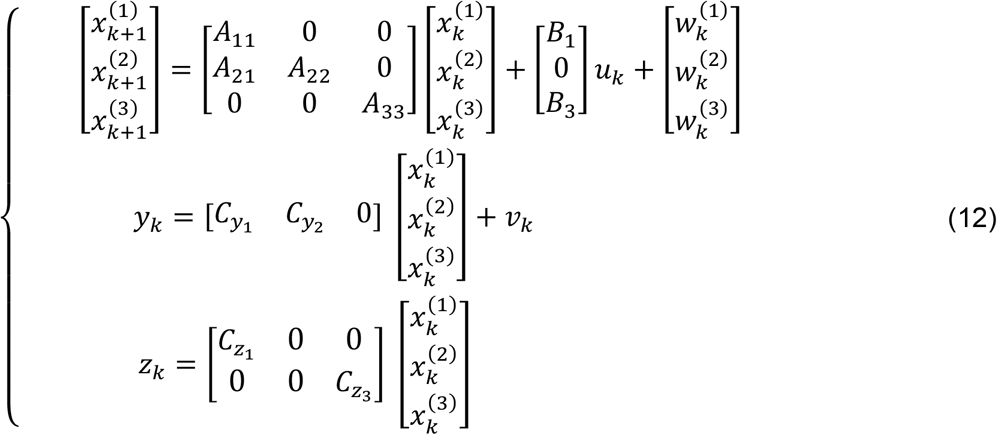

with

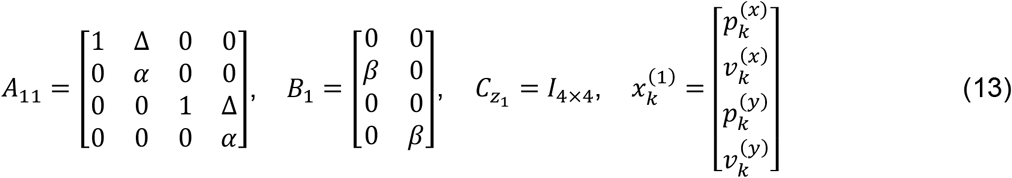

where Δ= 0.01 second is the duration of each time step, *α* = 0.99 is the damping ratio for velocity, and *β* = 0.01. The last two dimensions of behavior *z_k_* are only driven by states 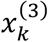 that are not reflected in the neural activity and are not engaged in the task, i.e. are not instructed to go to any target (similar to the vertical kinematics of finger position in real neural dataset 1, see ref. ^45^, because while the kinematics evolve in 3D, only 2 dimensions are used for control on a 2D plane). The OFC component of the brain controls the 2D cursor kinematics, which are the first 4 dimensions of *z_k_*, by generating an internal (unobservable) control command time series, which we will henceforth refer to as *c_k_*. The OFC implements a linear quadratic regulator (LQR). Briefly, let’s denote by 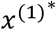 a desired target state for the behaviorally relevant state (i.e., target as dictated by task instructions). LQR determines the optimal online command *c_k_* as a linear function of the difference between the current state and the desired target state as

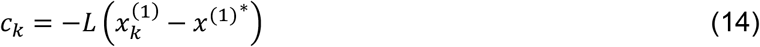

where 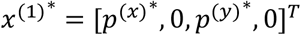, with 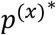 and 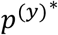 specifying the current target position according to task instructions, and the optimal feedback gain matrix *L* is obtained by solving the discrete algebraic Riccati equation^86^. Replacing the feedback equation (14) into equations (12) and (13) gives the full brain model including both the LSSM and the OFC components. Note that the overall measurable external input to the brain is the task instruction 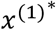, which we henceforth refer to as *u_k_*. Moreover, based on this full brain model, the behaviorally relevant block of the state transition matrix for the full brain model can be written as 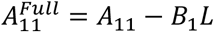. Thus, we compute the eigenvalues of 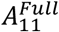 as the ground truth for the behaviorally relevant eigenvalues (**Fig. 5b**).

To generate a random realization of the simulated brain performing the task, we randomly choose a target among the permissible targets in the task (**Fig. 5c**). Then we iterate through equations (12) and (14), starting from the initial value *x*_-1_ = 0, until the brain state 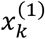 reaches the desired target 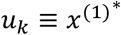 and stays within its boundary for 0.1 s (**Fig. 5c**). We then randomly choose a new target, update *u_k_* accordingly, and continue iterating through equations (12) and (14). Any time the desired target is reached, a new target is chosen and the data generation process continues as before. We generate *N* = 2000000 data samples with this procedure to get {*y_k_*,*z_k_*: 0 ≤ *k* < *N*} from equation (12). We also keep a timeseries of the desired target position values to use as the overall input 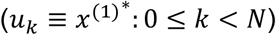 to the brain in IPSID and INDM models. Using cross-validation, we identify the model of the brain using IPSID, INDM, PSID and NDM.

#### Modeling non-human primate neural population activity during non-stereotypical movements

We study two publicly available neural datasets with distinct motor tasks recorded from three macaque monkeys. As the first dataset, we use primary motor cortex (M1) neural data from the Sabes lab^45^. In this experiment, the monkeys (monkey I and monkey L) performed continuous, self-paced reaches to targets chosen randomly with uniform probability from an 8×8 or 8×17 grid, without any time gaps or pre-movement intervals (**Fig. 6a**). The cursor was controlled based on the 2D position of the monkey’s fingertip in the horizonal plane. The task interface was presented to the monkey in a virtual reality environment. We analyze the first spike dimension available for each channel—resulting in 89 to 92 units for monkey I and 91 to 96 units for monkey L—from the first 7 recording sessions for each subject. For faster computation in our analyses, we randomly partitioned the units into two non-overlapping sets of equal sizes and analyzed each set separately. We compute the fingertip’s 2D velocity by taking derivative from the recorded 2D position. We take the measured 2D position as well as computed 2D velocity of the fingertip 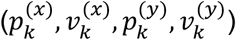 as the behavior signal *z_k_*.

As the second dataset, we use the dorsal premotor cortex (PMd) data recorded and made publicly available by the Miller lab^46,47^. In this experiment, the monkey (monkey T) performed sequential reaches to random targets on a screen by controlling a cursor via a two-link planar manipulandum (**Fig. 7a**). In this task, an on-screen visual cue (2 cm × 2 cm square) specified the target location for each reach and the monkey was given a liquid reward after making a series of four successful reaches. The location of each target was randomly chosen within 5-10 cm of the previous target. We analyzed single unit spiking activity during all 3 behavioral sessions, with 49, 46, and 57 single units. We take the measured 2D position and velocity of the arm 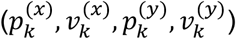 as the behavior signal *z_k_*.

For all three subjects, we use spike counts counted within non-overlapping 50 ms time intervals and smoothed by Gaussian kernel with a 50 ms s.d.^3,13,39,48^ as the neural activity *y_k_* (**Figs. 6, 7, S11**). Smoothing is performed as is typical in the field^3,13,39,48^. Gaussian distributed variables are commonly used to model both local field potentials (LFP)^14,19,30,44,61,65,66^ and spike counts^7,19,67,68^. We take the 2D target location in the task as the input time series (*u_k_*) provided to the subject. We perform all analyses within a 5-fold cross-validation and report the cross-validated CC of predicting behavior and neural activity as the performance measures (i.e., decoding and neural self-prediction).

To choose a suitable latent state dimension for our modeling of the intrinsic behaviorally relevant neural dynamics (**Figs. 6, 7, S11**), we estimate the latent state dimension that is sufficient for capturing most of the behavioral dynamics. To do this, we model the behavior time series using INDM as

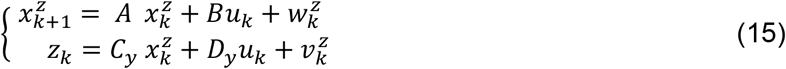

with different latent state dimensions in the range 1 ≤ *n_x_* ≤ 50 (**Fig. S13**). We then find the smallest latent state dimension for which the cross-validated one-step-ahead self-prediction of behavior reaches within 0.9 of its peak value. This dimension is found to be *n_x_* = 4 in all three subjects (**Fig. S13**). We use a horizon parameter of *i* = 40 (**Note S1**) for all (I)PSID and (I)NDM modeling in real neural datasets.

As a measure of the learned dynamics, we report the distribution of the learned eigenvalues in each subject by aggregating eigenvalues of the learned models across recording sessions and cross-validation folds, and adding up gaussian kernels (with s.d. of 0.05) centered around each learned eigenvalue. The mean of all gaussian kernels, when normalized to sum up to one over the unit disk, gives a probability mass function that estimates the probability of an eigenvalue being identified at each location on the complex plane in that subject (**Figs. 6b**, **7b, S11b**). We find the mode of the eigenvalue distributions, i.e., locations on the complex plane that have peak eigenvalue identification probability, for a given method and report their associated frequency and decay rate. The frequency and decay rate describe how a complex conjugate pair of eigenvalues at a given location on the complex plane would respond to an impulse excitation, i.e., they specify the ringing frequency of the response and how fast it would decay to less than 1% of its initial value. For a point *λ* = |*λ*|*e^jω^*, the associated frequency is computed as 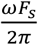 where *F_s_* = 20 *Hz* is the data sampling rate, and the decay rate is computed as 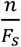 where *n* is the smallest integer for which |*λ*|^*n*^ < *e*^−1^.

To quantify how close the identified eigenvalues are to the input eigenvalues (**Figs. 6c**, **7c, S11c**), we compute the pointwise *KL-divergence* between the probability mass function found using the identified method *P_method_*, and the probability mass function of input eigenvalues *P_input_* as *D_KL_*(*P_input_* || *P_method_*). Similarly, to quantify the distance between the eigenvalue distributions found using a method for two subjects *i* and *j* (*P_subject_i__* and *P_subject_j__*), we computed their correlation coefficient, *symmetric KL-divergence*, defined as: 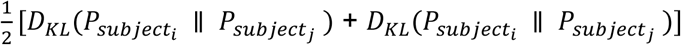, as well as the *mode distance* defined as the Euclidean distance between the locations with peak eigenvalue occurrence probabilities in *P_subject_i__* and *P_subject_j__*.

#### Statistics

We use the Wilcoxon signed-rank tests for all paired statistical tests.

### Supplementary Notes

#### Note S1 Preferential subspace identification in presence of inputs (IPSID)

##### Definitions

To simplify the description of IPSID, we define some notations. First, we define the notation

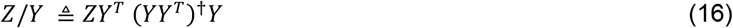

to denote the orthogonal projection^40^ of the wide matrix 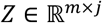 onto another wide matrix 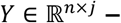 where the superscript † denotes the pseudoinverse operation. An orthogonal projection can also be thought of as the linear minimum mean squared prediction of columns of *Z* using columns of *Y*. This is because *ZY^T^*(*YY^T^*)^†^ is equal to cross-covariance of columns of *Z* and *Y*, multiplied by the inverse of the covariance of the columns of *Y*. Second, we define the shorthand notation^40^

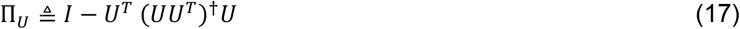

where 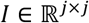 is the identity matrix. Π_*u*_ is a matrix that when multiplied from the left by a matrix *Z*, would remove the orthogonal projection of *Z* onto *U* from *Z*. In other words, we have 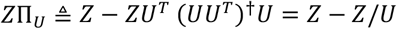 for any matrix 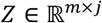. Third, we define the notation

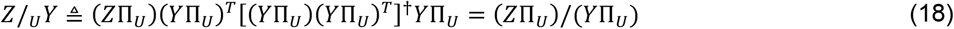

to denote the oblique (i.e. non-orthogonal) projection^40^ of matrix *Z* onto *Y* along matrix 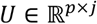. Intuitively, the oblique projection in equation (18) means that we first find the part of *Z* that is not predictable from *U* (i.e. *Z*Π_*U*_), and then project that part onto the part of *Y* that is not predictable from *U* (i.e. *Y*Π_*U*_). This gives us the part of *Z* that is predictable by *Y* but not explained by *U*.

We also define the following matrices form the training neural time series 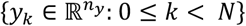

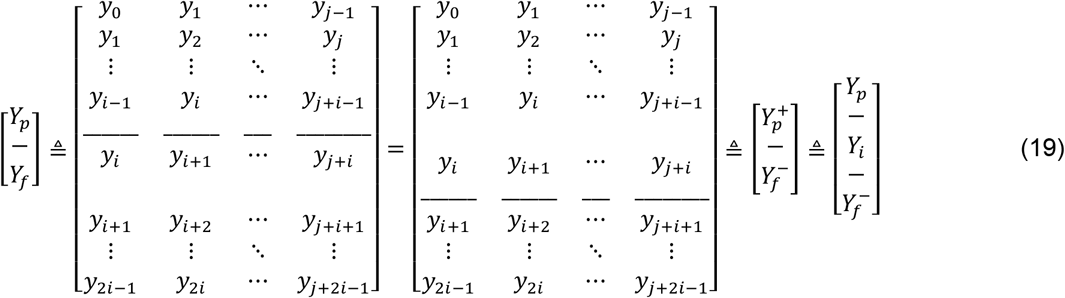

and analogously define the following from the training behavior time series 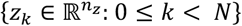

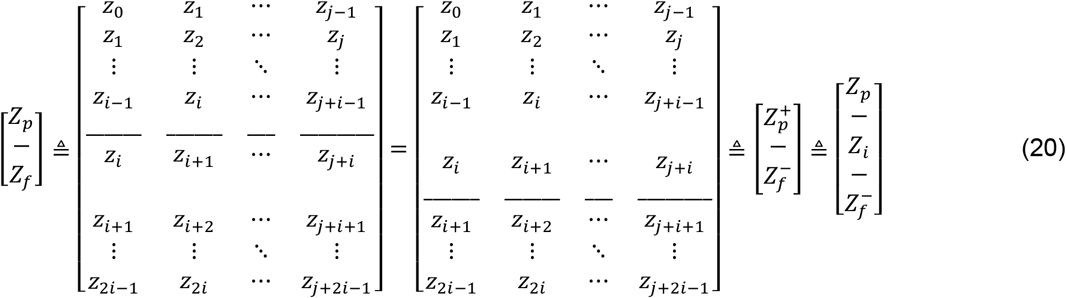

and the following from the input time series 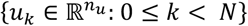

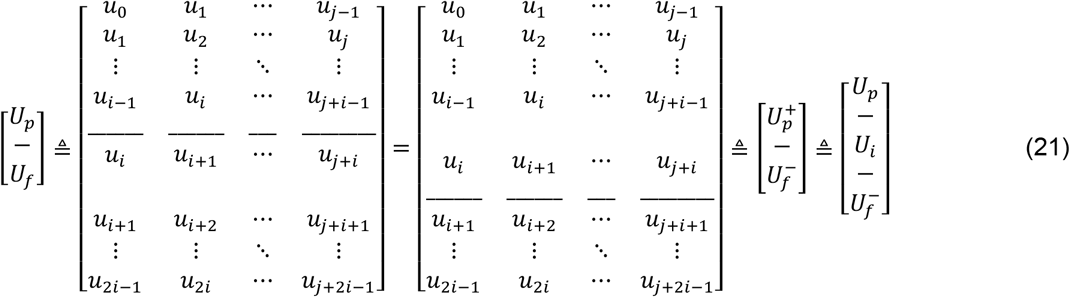

where *i* is a provided quantity termed the projection horizon, and *j* is number of columns in the abovementioned block matrices. Given the number of data samples *N*, we set *j* = *N* – 2*i* to use all data samples.

##### Outline of the IPSID algorithm

Here, we describe the IPSID algorithm. Given the neural, behavioral, and input training time series, i.e. 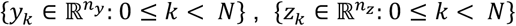 and 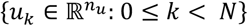, respectively, and given the state dimension *n_x_*, parameter *n*_1_ ≤ *n_x_*, and projection horizon *i*, IPSID learns the parameters of the model in equation (1) while prioritizing the learning of intrinsic behaviorally relevant states. Doing so, IPSID can dissociate intrinsic behaviorally relevant neural dynamics from input dynamics and from other intrinsic neural dynamics.

##### Stage 1: Extract *n*_1_ latent states directly from data via an oblique projection of future behavior onto past neural activity and past input, along future inputs

1. Form examples of future behavior *Z_f_* (equation (20)) and the associated past neural activity *Y_p_* (equation (19)). Also form the corresponding samples of future and past input *U_f_* and *U_p_* (equation (21)). Project *Z_f_* onto *Y_p_* and *U_p_* along *U_f_* to get

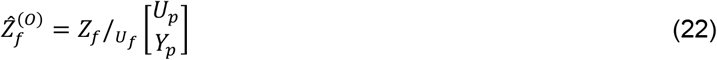

where oblique projection is defined as in equation (18).
2. Compute the singular value decomposition (SVD) of 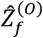 and keep the top *n*_1_ singular values:

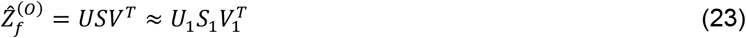
3. Compute the intrinsic behaviorally relevant latent state as

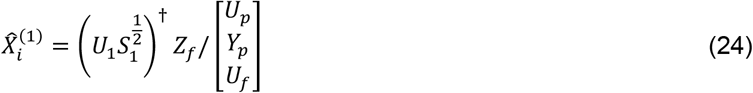

##### Stage 2 (optional): extract *n_x_* – *n*_1_ additional latent states via an oblique projection of residual future neural activity onto past neural activity and past input, along future input

1. Find the prediction of *Y_f_* from 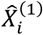 and subtract this prediction from *Y_f_* using an oblique projection to keep the part that is predictable from *U_f_* and *U_p_*. Name the result 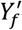 (i.e., residual future neural activity):

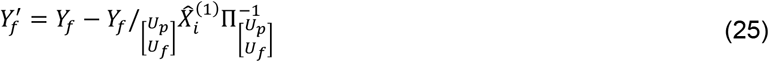 Here 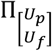 is defined per equation (17).
2. Project the residual future neural activity 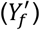 onto *Y_p_* and *U_p_* along *U_f_* to get

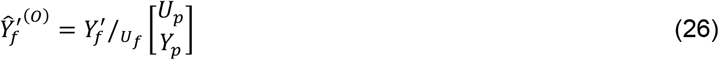
3. Compute the SVD of 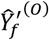, and keep the top *n_x_* – *n*_1_ singular values:

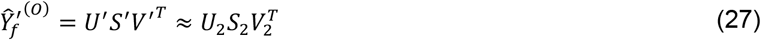
4. Compute the remaining latent states as

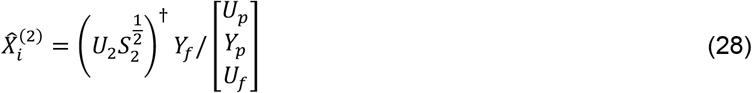

##### Final step: given the extracted latent states, identify model parameters

1. If stage 2 is used, concatenate 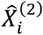 to 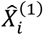 to get the full latent state 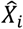, otherwise take 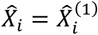.
2. Repeat all steps with a shift of one step in time to extract the states at the next time step 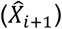. To shift the time step, use 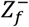 (equation (20)), 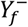 (equation (19)), 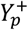 (equation (19)), 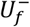 (equation (21)), and 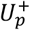 (equation (21)) instead of *Z_f_*, *Y_f_*, *Y_p_*, *U_f_* and *U_p_*, respectively.
3. Compute *A*_11_, *A*_21_, *A*_22_, *C_y_* and *C_z_* based on least squares solutions of equations (6) as

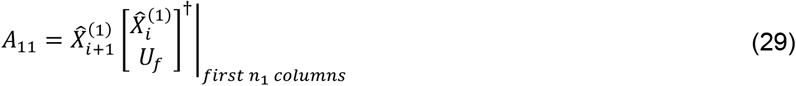

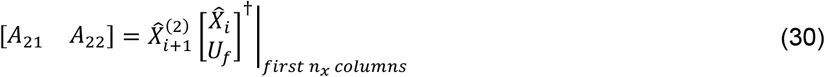

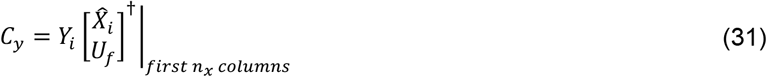

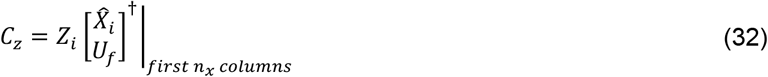

where *Y_i_* and *Z_i_* are as defined in equations (19) and (20), respectively.
4. Compute an estimate of the noise time series *w_k_* and *v_k_* from equation (1) based on the residuals/errors of the least squares solutions from the previous step as

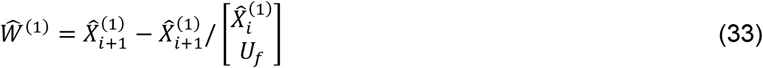

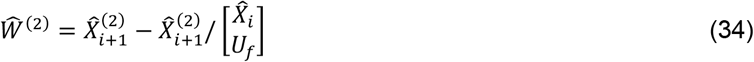

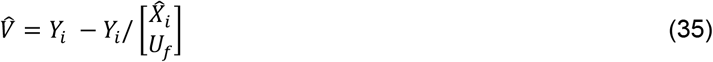
5. Compute the covariances and cross-covariance of *w_k_* and *v_k_* as

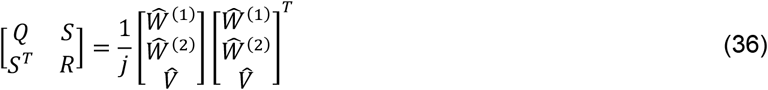
6. Follow a procedure similar to ref. ^40^, pages 125-127, to find the least squares solution for the model parameters *B* and *D_y_*.
7. This concludes the learning of all model parameters in the first two rows of equation (1) and thus we can now run a Kalman filter to recursively estimate the latent states (without looking at behavior) as

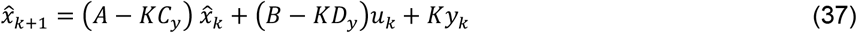

where *K* is the steady state Kalman gain (**SI Methods** equation (7)).
8. Finally, compute the parameter *D_z_*, which captures the direct non-dynamic effect of input on behavior, via an orthogonal projection as

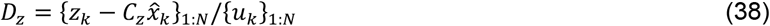

where {*α_k_*}_1:*N*_ denotes constructing an *N*-column matrix with column *n* containing *α_n_*.

The above concludes the learning of all model parameters using IPSID. For the special case of *n*_1_ = 0, only stage 2 will be performed and the algorithm will reduce to INDM^40^, which does not prioritize or dissociate the learning of intrinsic behaviorally relevant neural dynamics.

#### Note S2 IPSID for scenarios where neural recordings do not capture all downstream regions of input that influence behavior

Here, we describe how IPSID can also support scenarios where the recorded neural activity does not cover all of the downstream regions of the input. In other words, as opposed to the main IPSID algorithm (**Note S1**), here we allow existence of latent dynamics that are affected by input and contribute to the generation of behavior but are not encoded in the recorded neural activity (i.e., 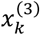 in equation (2)). We now outline the key steps that need to be performed in addition to the steps described in **Note S1** to enable this extension. Compared with **Note S1**, here in addition to the state dimension *n_x_* and the parameter *n*_1_ specifying the dimension of the behaviorally relevant latent states, an additional parameter *n*_2_ should also be specified by the user that determines the dimension of 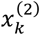, i.e., the intrinsic latent states extracted beyond those that are behaviorally relevant. In **Note S1**, *n*_2_ was automatically inferred as *n*_2_ = *n_x_* – *n*_1_; but here the user can specify any *n*_2_ in the range 0 ≤ *n*_2_ ≤ *n_x_* – *n*_1_. This version of the IPSID algorithm then learns the parameters of the model in equation (2), where a new set of states denoted by 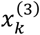 with *n_x_* – *n*_1_ – *n*_2_ dimensions describe the behavior dynamics that are predictable from input but are not reflected in recorded neural activity.

##### Initial projection step

Find the part of future behavior that is encoded in neural activity.

1. Model the neural signal *y_k_* using only stage 2 of IPSID (i.e., INDM) to extract *m* latent states 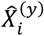 that describe the neural activity, where *m* is the total number of latent states that drive the neural activity (sum of dimensions of 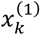 and 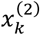 in the true model of equation (2)). The appropriate value of *m* for a given data can be correctly estimated by increasing *n_x_* in the model until neural self-prediction reaches a peak performance (**Fig. S8b**). For analysis in real data (**Figs. 6, 7, S11**), we use *m* = 150.
2. Form examples of future behavior *Z_f_* (equation (20)) and find its oblique projection onto 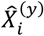 along *U_f_* and *U_p_*, naming the result 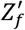

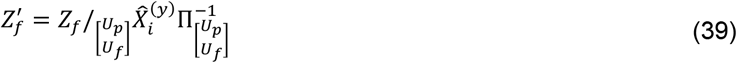

where oblique projection is defined as in equation (18) and 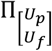 is defined as in equation (17).

The above steps extract the part of the future behavior that is encoded in the recorded neural activity, i.e., 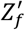. We next perform IPSID stages 1 and 2 as in **Note S1**, with the only difference being the use of 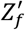 instead of *Z_f_*.

***Stage 1:** perform the IPSID stage 1 as outlined in **Note S1** but use* 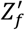 (*the part of future behavior that is encoded in neural activity) instead of Z_f_, to extract n*_1_ *intrinsic behaviorally relevant neural states* 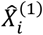.

***Stage 2 (optional):** perform the IPSID stage 2 as outlined in **Note S1**, to extract **n***_**2**_ *additional neural latent states* 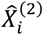.

##### Parameter learning step for neural dynamics

given the extracted neural latent states, learn all model parameters in equation (2) that are related to the neural dynamics.

1. Apply the final parameter learning step of **Note S1** to learn all model parameters in equation (2) that are associated with the recorded neural dynamics i.e., *A*_11_, *A*_21_, *A*_22_, *B*_1_, *B*_2_, 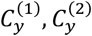, *D_y_* and noise statistics.
2. Learn the 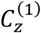 parameter from equation (2) as

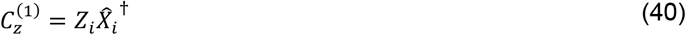

where *Z_i_* is as defined as in equation (20).

This concludes the learning of all intrinsic neural dynamics. We next proceed with an optional step that if desired can learn additional behavior dynamics that are predictable from input but are not encoded in neural activity (i.e., 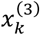 in equation (2)).

##### Learn input-driven behavior dynamics not encoded in recorded neural activity (Optional)

*extract n_x_ – n*_1_ – *n*_2_ *additional latent states that are not encoded in neural activity but describe behavior dynamics predictable from input*.

1. Run the Kalman filter for the learned neural model (equation (7)) to estimate neural latent states 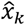 and the prediction of behavior using those latent states (equation (8)). Then subtract this prediction from the behavior signal to get the residual behavior signal 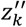 that is not predictable from the neural latent states.
2. Apply IPSID stage 2 (i.e. INDM) to 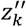 instead of *y_k_* to extract *n_x_* – *n*_1_ – *n*_2_ latent states 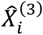 and learn their associated model parameters *A*_33_, *B*_3_, and 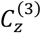 from equation (2) to conclude this version of IPSID.

### Supplementary Figures

**Fig. S1.**
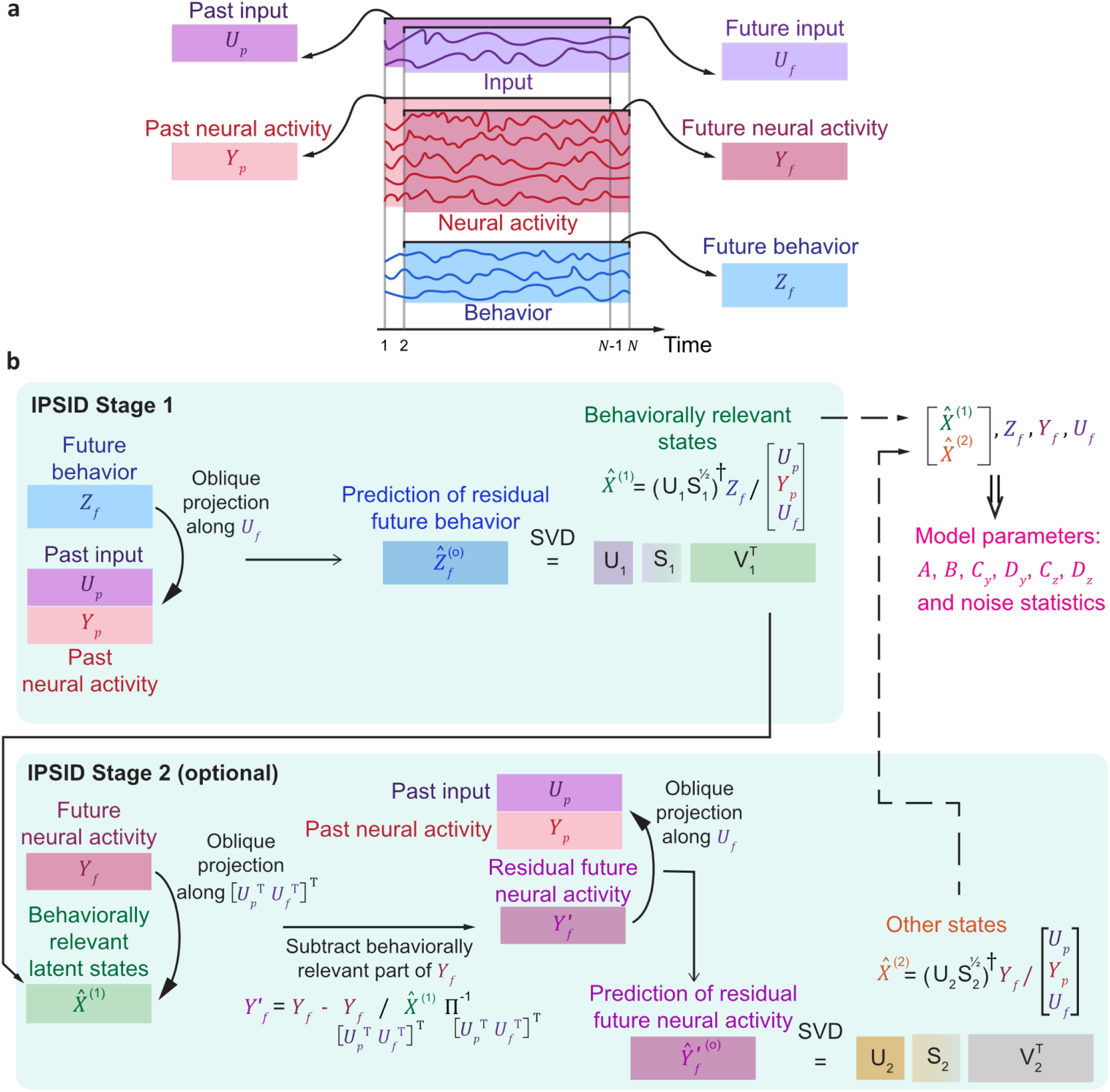
Visualization of the IPSID algorithm. **(a)** The extraction of future and past training data is shown. Here, *U*, *Y* and *Z* denote input, neural activity and behavior, respectively. Colored rectangles represent data matrices used to extract the latent state (see **Note S1** for general definition). Future matrices *U_f_*, *Y_f_* and *Z_f_* are constructed by shifting the columns of the past matrices one step ahead in time. **(b)** In the first stage of IPSID, the intrinsic behaviorally relevant states 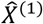 are extracted from data with priority with the following procedure: Future behavior *Z_f_* is projected onto the concatenation of past input *U_p_* and past neural activity *Y_p_* along the subspace spanned by the future input *U_f_* to obtain 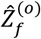, which is the prediction of residual future behavior where residual refers to the part not predictable by future inputs (**Note S1**). Performing SVD on 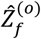 gives the intrinsic behaviorally relevant states 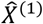. In the second stage of IPSID, which is optional, any remaining intrinsic latent states 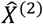 are extracted from the data with the following procedure: The predictable part of future neural activity *Y_f_* from 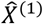 is removed to obtain the residual future neural activity 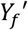, i.e., the part not predictable by 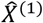 (operator Π is defined in **Note S1**). Further, oblique projection of 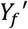 onto concatenation of the past input and past neural activity along the subspace spanned by the future input *U_f_* results in 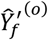, which is the prediction of residual future neural activity, where residual refers to the part not predictable by future inputs or by 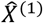. Performing SVD on 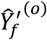 gives the remaining intrinsic latent states 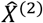. Once the latent states 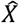 are extracted—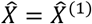 if only using the first stage or 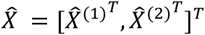 if also using the optional second stage—, model parameters can be learned using linear regression (**Note S1**).

**Fig. S2.**
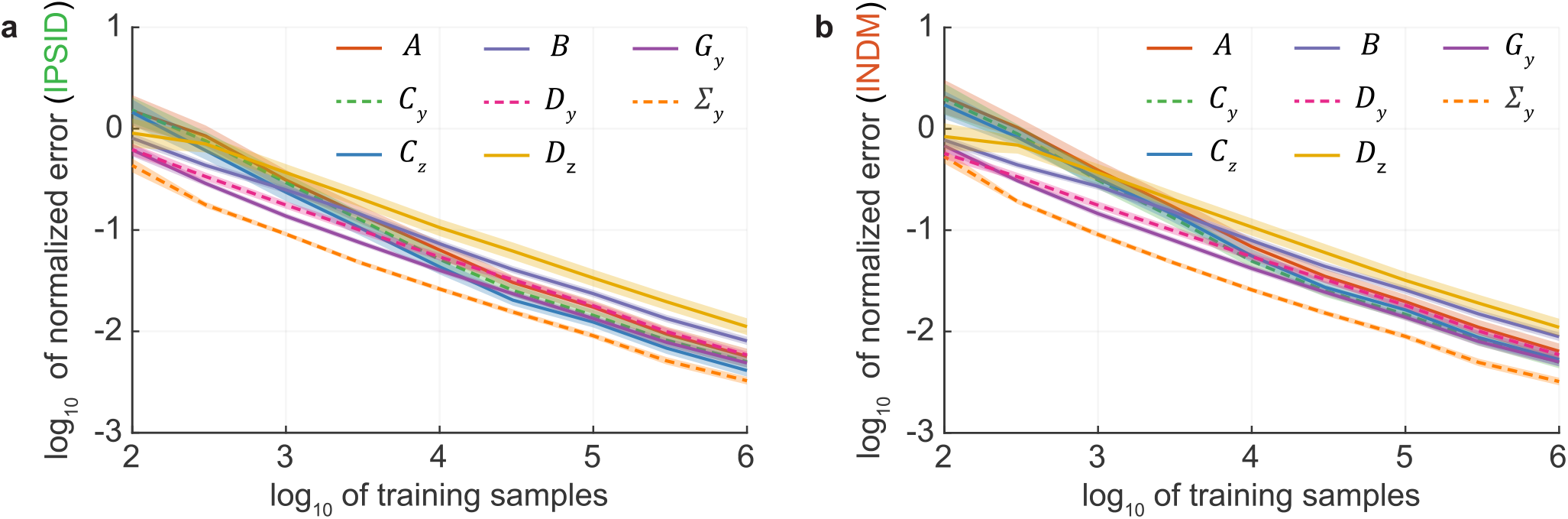
IPSID correctly learns all model parameters. **(a)** Normalized error in learning each model parameter using IPSID versus the number of training samples used for learning. Models are as in equation (1). Solid lines show the mean across the random models and the shaded areas show the s.e.m. (*n* = 100 random models). Parameters 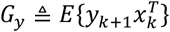 and 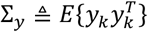 fully represent the noise statistics, unlike *Q*, *R*, and *S* from equation (5) in **SI Methods**, which are a redundant representation that is not uniquely identifiable and thus is not suitable for evaluating model identification methods^19,40,80^. **(b)** same as (a) for INDM.

**Fig. S3.**
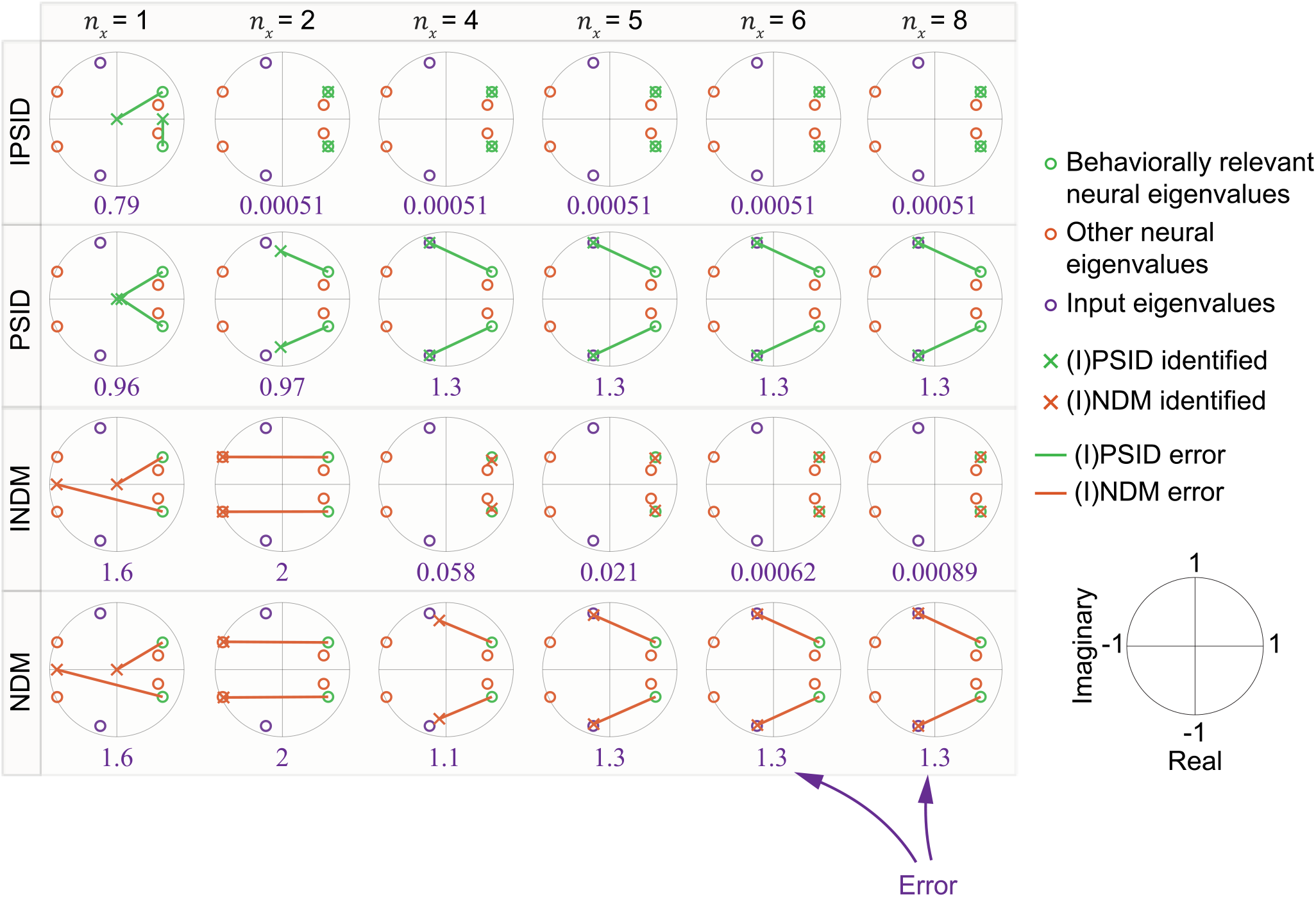
Unlike other methods, IPSID correctly learns the intrinsic behaviorally relevant neural dynamics in the presence of input even when using lower-dimensional latent states (i.e., even when performing dimensionality reduction). For one simulated model as in equation (1), the identified intrinsic behaviorally relevant eigenvalues of the state transition matrix *A* are shown for (I)PSID and (I)NDM for different latent state dimensions. These eigenvalues quantify the dynamics (**SI Methods**). True model eigenvalues are shown as colored circles, with colors indicating their relevance to input, neural activity, behavior, or both. Crosses show the identified behaviorally relevant eigenvalues when modeling the neural activity. When the state dimension *n_x_* is less than the true dimension of behaviorally relevant states (*n*_1_ = 2), missing eigenvalues are taken as 0, representing an equivalent model for which *n*_1_ – *n_x_* latent state dimensions are always 0. Thus, all cases have 2 crosses indicating 2 identified eigenvalues (*n*_1_ – *n_x_* of which are zero when *n_x_* < *n*_1_). Lines indicate the error of the identified eigenvalues. The normalized value of the error—average line length normalized by the average true eigenvalue magnitude—is noted below each plot (**SI Methods**). Unlike IPSID, INDM may learn dynamics that are unrelated to behavior at lower state dimensions (i.e., when performing dimensionality reduction). NDM and PSID do not consider input and thus may learn dynamics that are confounded/influenced by input dynamics.

**Fig. S4.**
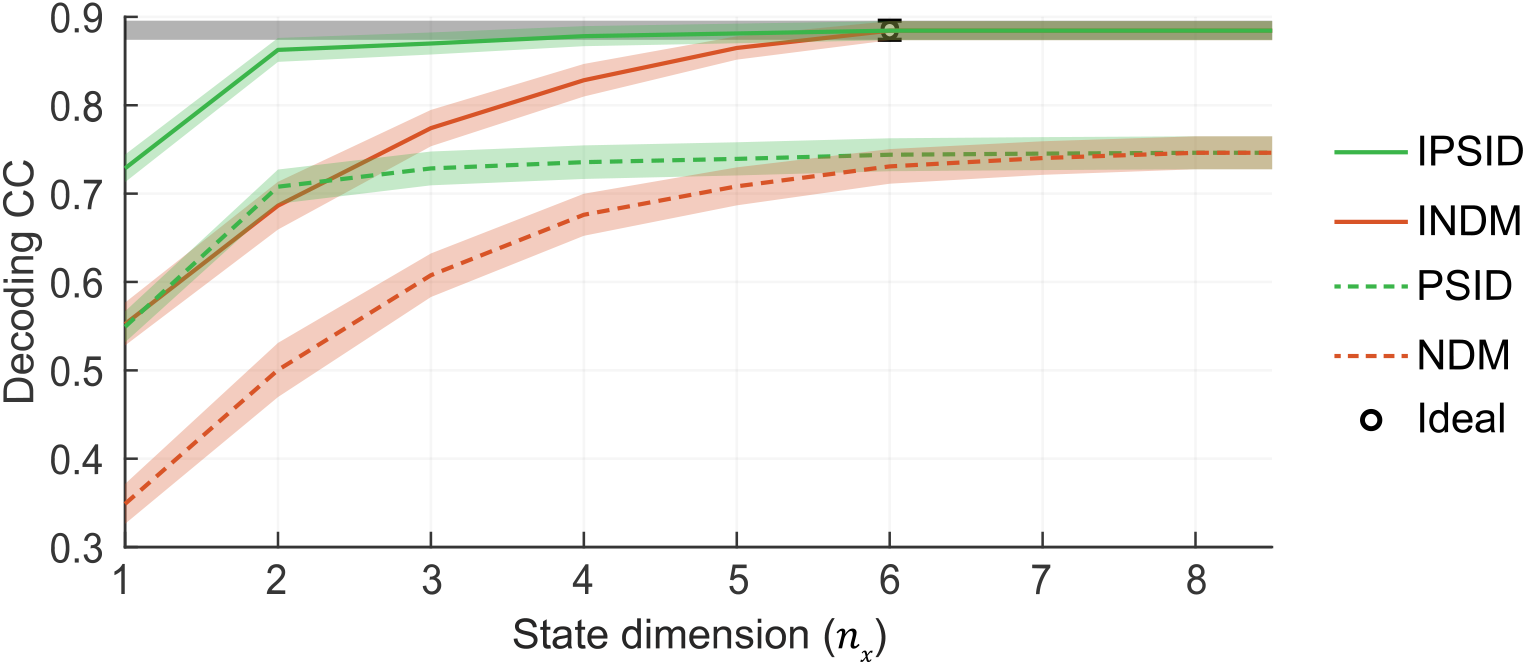
Quantified by behavior decoding accuracy, IPSID correctly prioritizes learning of intrinsic behaviorally relevant neural dynamics in the presence of input. Cross-validated behavior decoding correlation coefficient (CC) given the same random models as **Fig. 2b**. Ideal decoding CC using true model parameters is shown in black. IPSID/INDM consider input and thus are able to achieve peak decoding similar to the ideal decoding, with IPSID achieving the peak decoding using much lower-dimensional latent states than INDM. This shows the IPSID correctly prioritizes the learning of intrinsic behaviorally relevant neural dynamics unlike INDM. NDM/PSID do not consider input and thus do not reach ideal behavior decoding accuracy even with high-dimensional latent states. Note that for models that have input (i.e., those learned with IPSID and INDM), the input is observed for decoding and thus here we use behavior decoding accuracy simply as a measure of how well the behaviorally relevant neural dynamics are explained by the model and not as a measure of pure neural decoding of behavior.

**Fig. S5.**
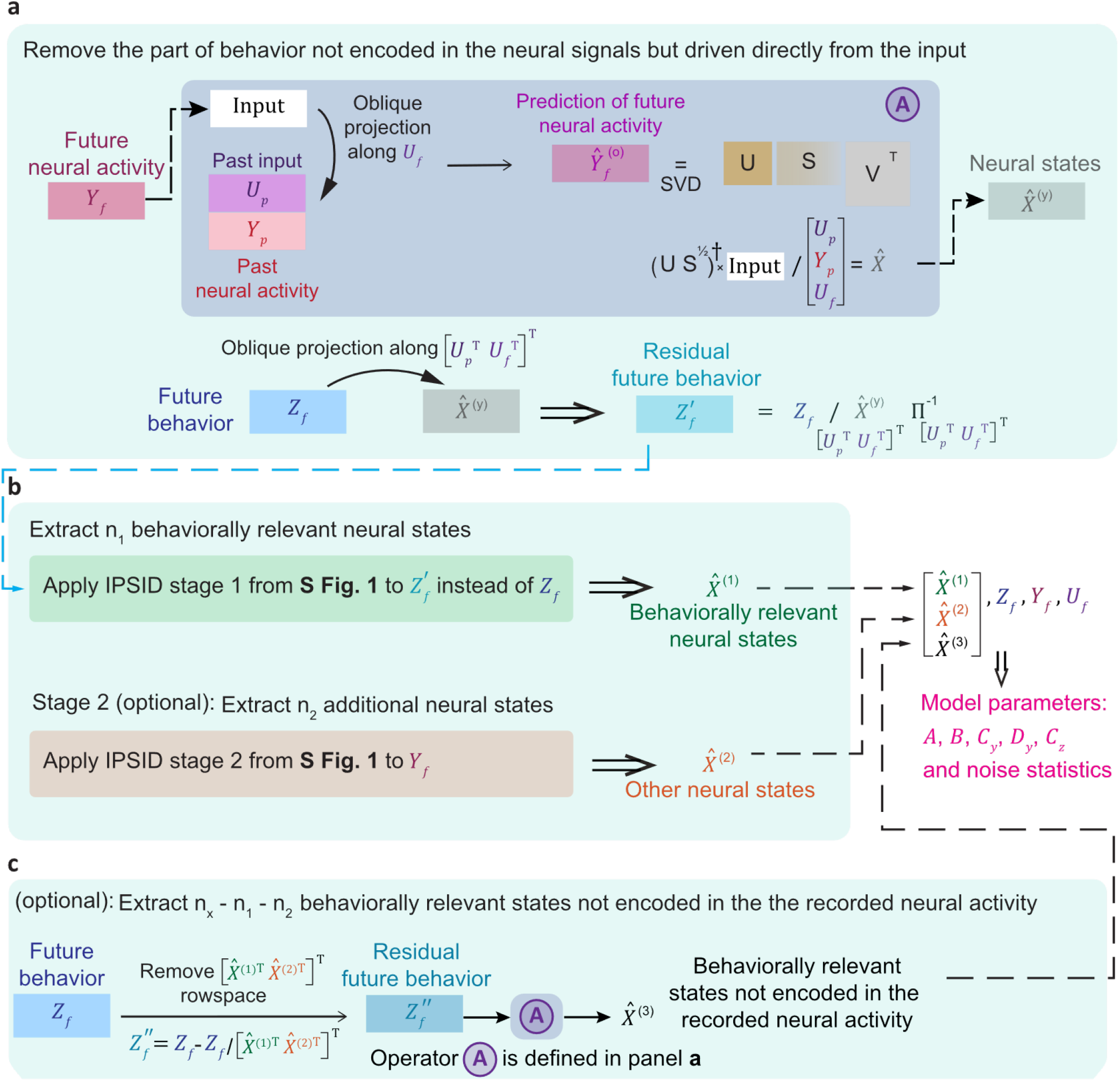
Visualization of IPSID algorithm with support for scenarios where the recorded regions do not cover all downstream regions of the input. **(a)** To support the scenario where recorded regions do not cover all downstream regions of input, we add an additional step that precedes the IPSID two-stage approach presented in **Fig. S1** and **Note S1**. Specifically, before extraction of intrinsic behaviorally relevant latent states in stage 1, the future behavior *Z_f_* is projected onto a latent state representation, termed 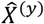, of all neural activity (operator Π is defined in **Note S1**); we extract the 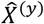 using the second stage of IPSID (the purple box which is also called operator 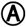). The dimension of the latent representation 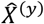 should be chosen high enough such that neural dynamics are well-represented in 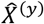. In analyses of real data, we determine this dimension by forming a plot of neural self-prediction using 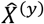 vs the dimension of 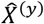, and finding a dimension that reaches a self-prediction close to the peak (**Fig. S8b**). We refer to the result of the projection of future behavior *Z_f_* onto the latent representation 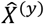 as 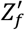, which is thus the part of future behavior dynamics that is reflected in the neural recordings. This step excludes any behavior dynamics that are not represented in the recorded neural activity from being learned in stage 1. **(b)** Next, IPSID stage 1 and 2 follow as explained in **Fig. S1** and **Note S1**, but this time the residual future behavior 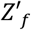 (from panel a) is used in these stages instead of the original future behavior *Z_f_* (see **Note S2)**. This concludes the learning of intrinsic behaviorally relevant neural dynamics (stage 1) and dissociating them from other intrinsic neural dynamics (stage 2). **(c)** Subsequently, we can optionally learn behavior dynamics that are predictable from input but are not reflected in the recorded neural activity. To do so, in a second optional step, we compute the residual behavior that is not yet predictable using the already-extracted latent states (i.e., using 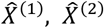), and apply the second stage of IPSID (operator 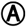) on that residual behavior signal. This is done by replacing future and past neural signals with future and past residual behavior signals in this second stage. This step allows us to build a model that predicts these residual behavior dynamics purely using the input via forward prediction. This concludes the dissociation of behavior dynamics that are encoded in the neural activity from those that are predictable from input but are not encoded in the recorded neural activity.

**Fig. S6.**
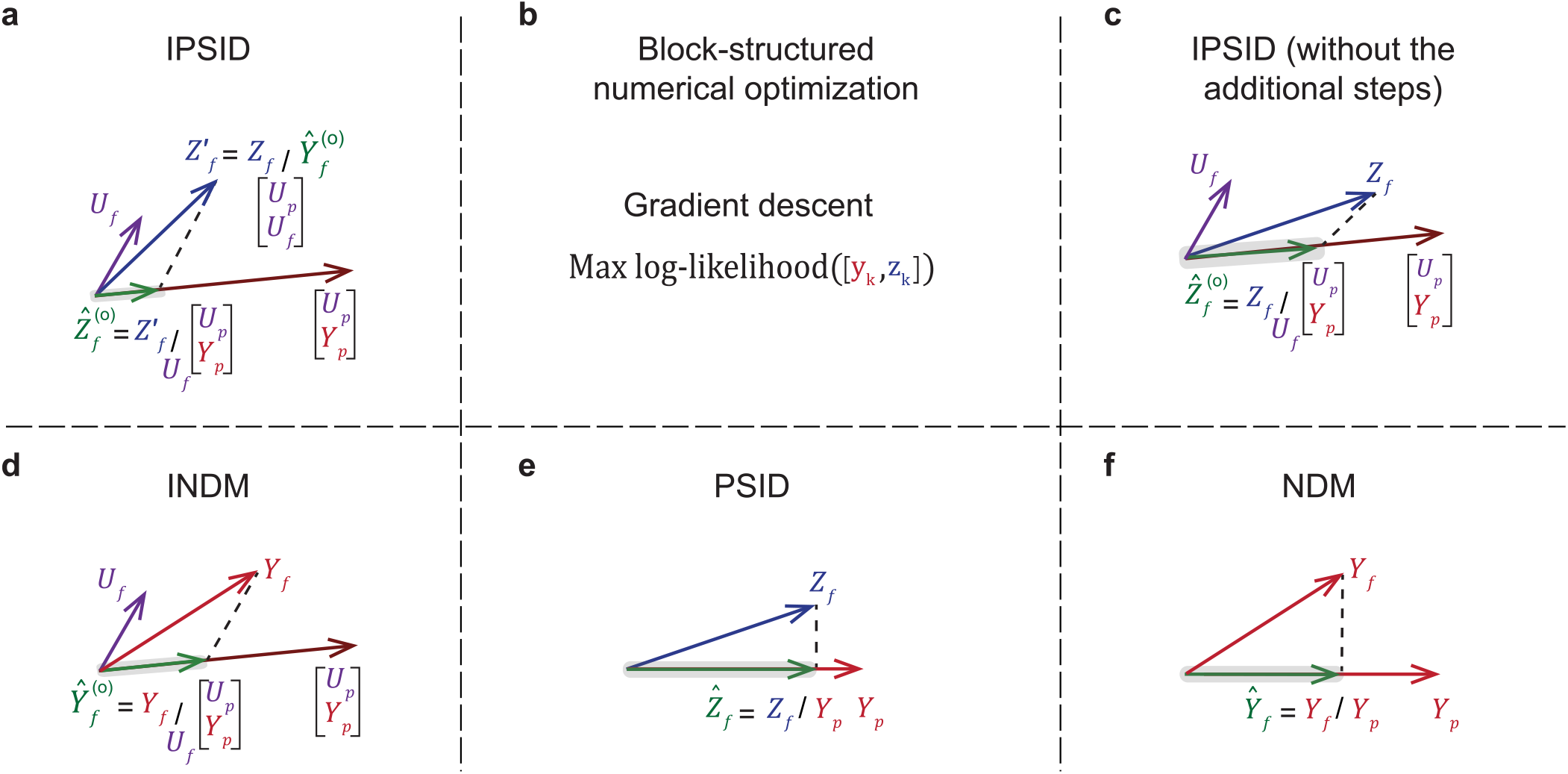
Simplified schematic overview of the key operations for subspace and numerical optimization learning methods. A schematic overview of the key operations involved in **(a)** IPSID, **(b)** block-structured numerical optimization, **(c)** IPSID (without additional steps), **(d)** INDM, **(e)** PSID and **(f)** NDM. *Z_f_*, *Y_f_*, *Y_p_*, and *U_f_*, *U_p_* denote future behavior, future and past neural activity, and future and past input, respectively (**Note S1**, **Fig. S1**). A/B denotes an orthogonal projection of A onto B and *A/_C_B* denotes an oblique projection of A onto B along C (**Note S1**). A block-structured numerical optimization method is additionally compared which learns the model parameters of equation (6) from **SI Methods** via gradient descent (**SI Methods**).

**Fig. S7.**
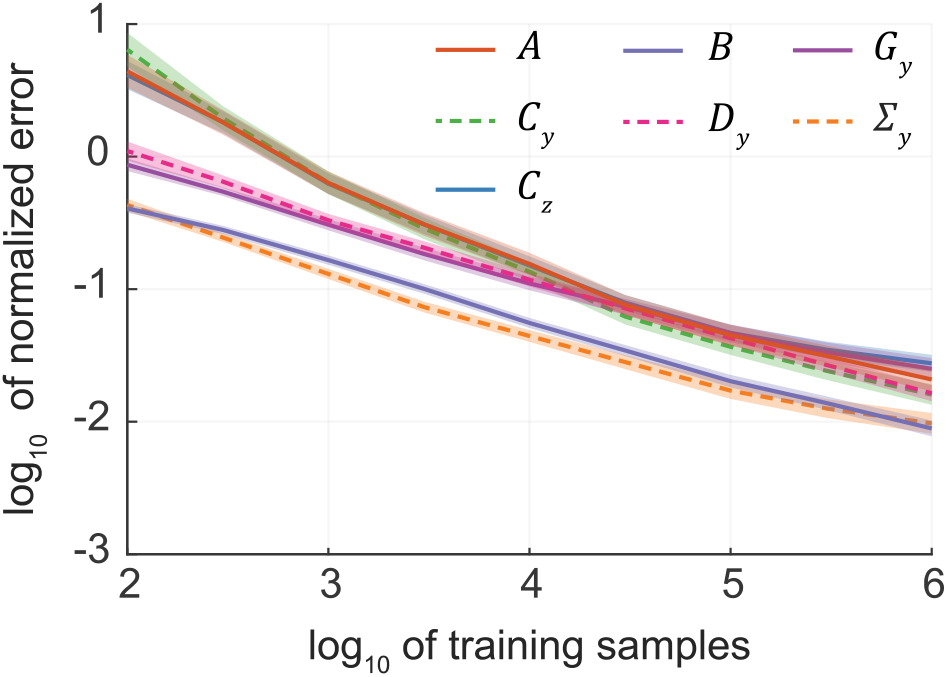
IPSID correctly learns model parameters even in scenarios where the recorded regions do not cover all downstream regions of the input. Notation is as in **Fig. S2a**, but for simulating models where some behavior dynamics that are influenced by inputs are not reflected in the recorded neural activity. Here 100 random models are simulated according to equation (2). Evaluated parameters are for the parts of the model that are relevant to the neural states in the recordings, i.e., 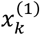 and 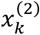. Learning 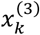 entails stage 2 of the same method as was validated in **Fig. S2a** (**Fig. S5c**); this 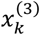 learning is also validated in a different way in **Fig. S8** next.

**Fig. S8.**
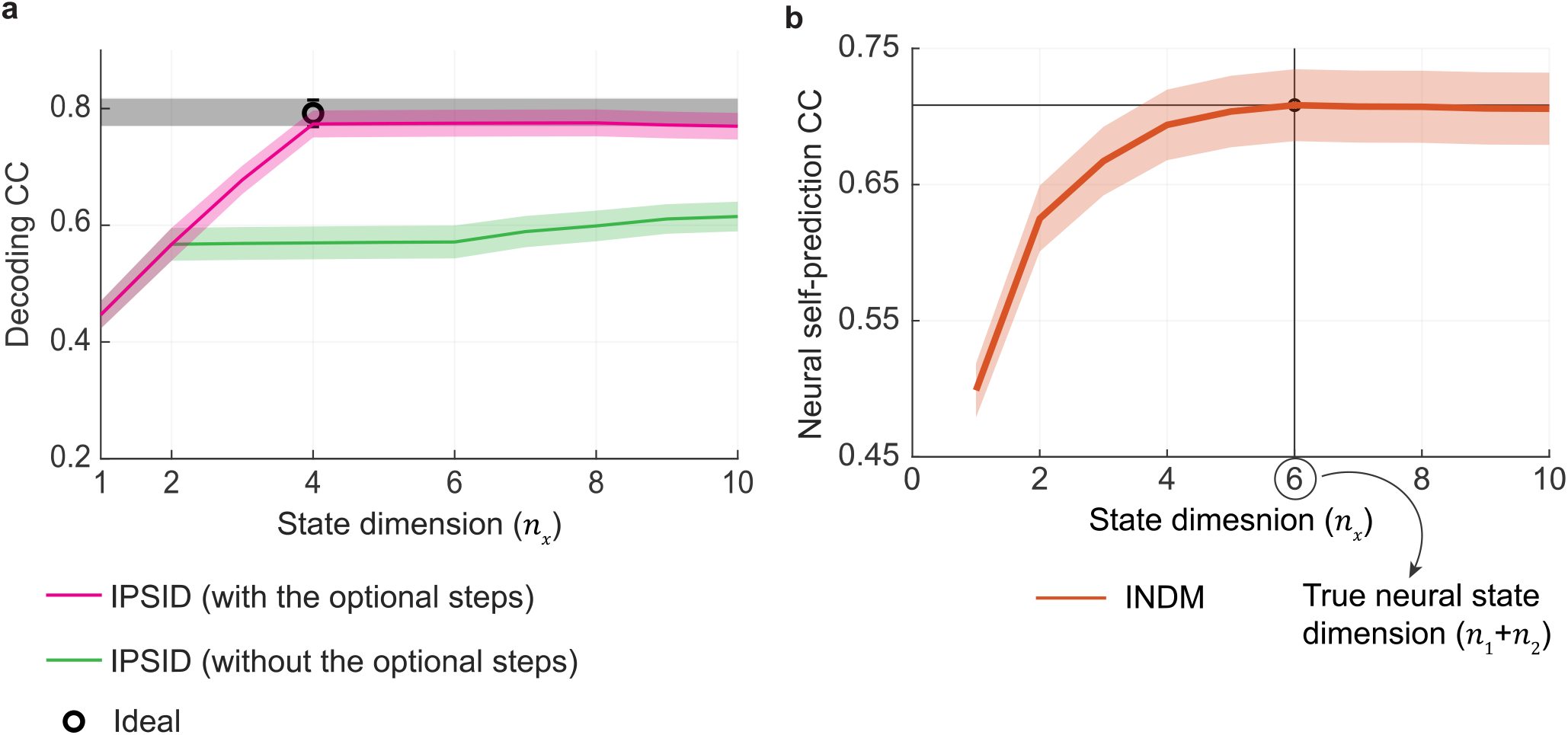
Quantified by behavior decoding accuracy, IPSID can achieve ideal prediction of behavior from input and neural activity even in scenarios where the recorded regions do not cover all downstream regions of the input. **(a)** Cross-validated behavior decoding correlation coefficient (CC) given 100 random models generated according to equation (2), where some latent states 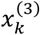 reflect the influence of input on behavior but are not reflected in recorded neural activity *y_k_*. Ideal behavior decoding CC using the true full model in equation (2), which also includes the 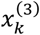 state components, is shown in black. IPSID can also optionally (magenta) learn and dissociate any dynamics in behavior that are predictable by input but are not encoded in recorded neural activity, i.e., 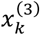. Thus, IPSID is able to achieve peak behavior decoding similar to the ideal model decoding (**Fig. S5c**, **Note S2**). **(b)** Neural self-prediction vs. state dimension for models learned using the second stage of IPSID alone. The latent state dimension that reaches peak self-prediction is equal to the true dimension of neural activity (i.e., *n*_1_ + *n*_2_). This procedure of finding the state dimension for peak neural self-prediction is thus used to determine the dimension of 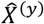 in the first optional step of IPSID (see **Fig. S5**). As shown here, this procedure correctly reveals the latent state dimension required for capturing the neural dynamics in 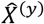 because the dimension to reach the peak neural self-prediction in (b) is equal to the true neural state dimension.

**Fig. S9.**
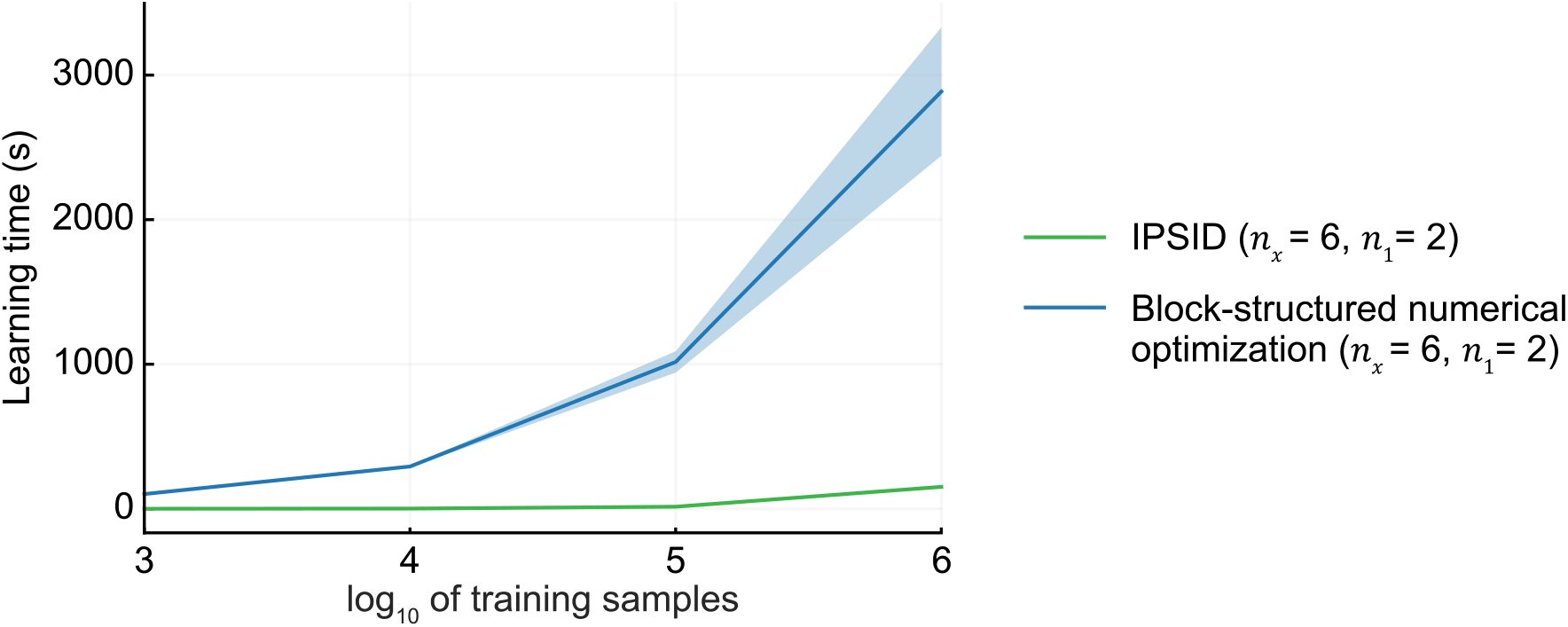
IPSID learns the model parameters faster than numerical optimization in Fig. 4. Learning time for IPSID and block-structured numerical optimization is shown as function of training samples. For all training sample sizes, IPSID is significantly faster than the numerical optimization for learning because it involves a pre-specified set of linear algebraic operations rather than iterative learning via gradient descent. Using different deep learning libraries and adjusting other implementation details could improve the speed of these methods. Nevertheless, because IPSID uses an analytical and non-iterative method, it would likely be generally faster in terms of learning speed compared with iterative numerical optimization approaches, but the exact comparison will depend on various implementation factors and the results here are just for our implementation described in **SI Methods**.

**Fig. S10.**
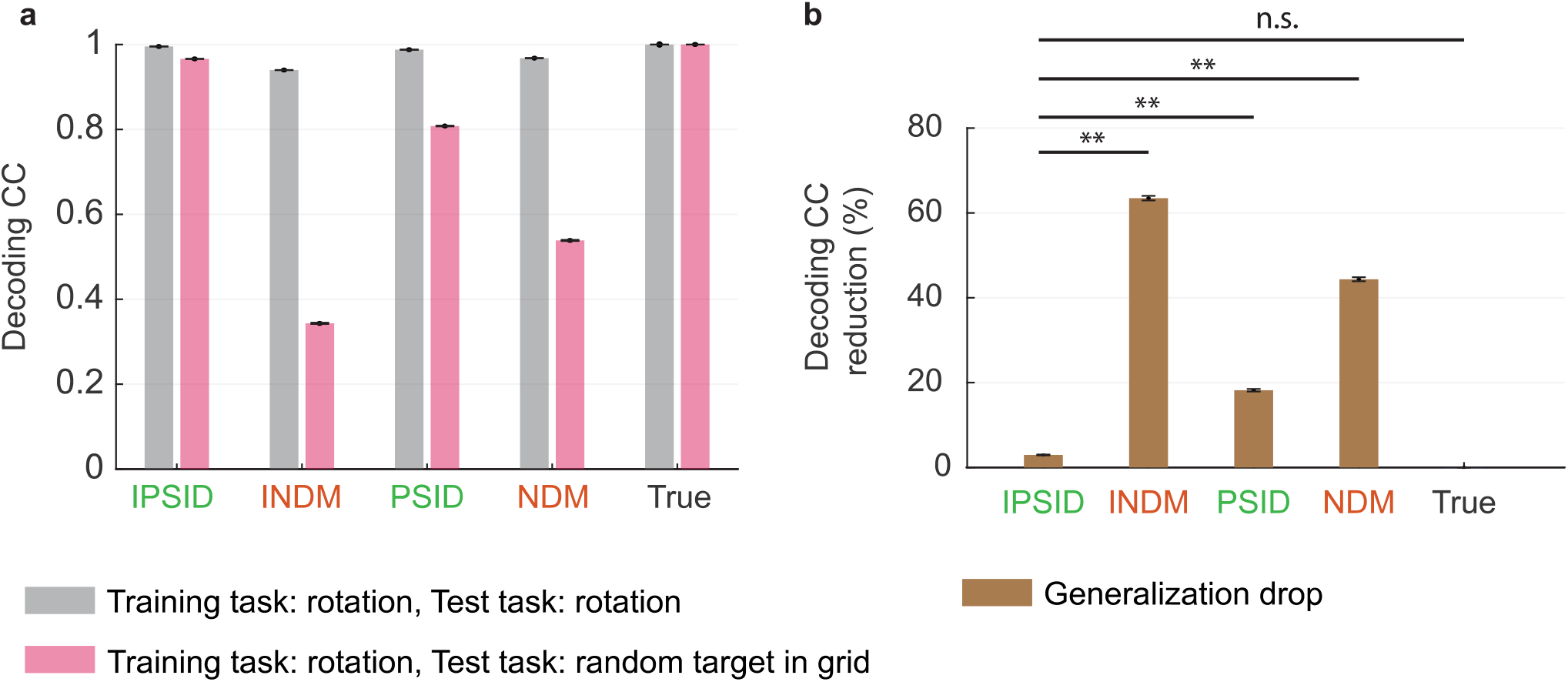
Models learned by IPSID generalized across behavioral tasks. Models were trained using data collected during a rotation task. **(a)** Performance is shown both for test data collected during the same rotation task used in training (task 1 from **Fig. 5**) and for test data collected during a different random target in grid task (task 3 from **Fig. 5**). Performance is averaged across 10 simulations of each task. Bars show the mean and whiskers, which are very short, show the s.e.m. **(b)** Relative drop (%) in behavior decoding correlation coefficient when generalizing a model trained on data from a rotation task to data from a random target in grid task. Double asterisks indicate *P* < 0.005 and n.s. indicates *P* > 0.05 for a one-sided signed-rank test. IPSID is the only method that generalizes well across tasks.

**Fig. S11.**
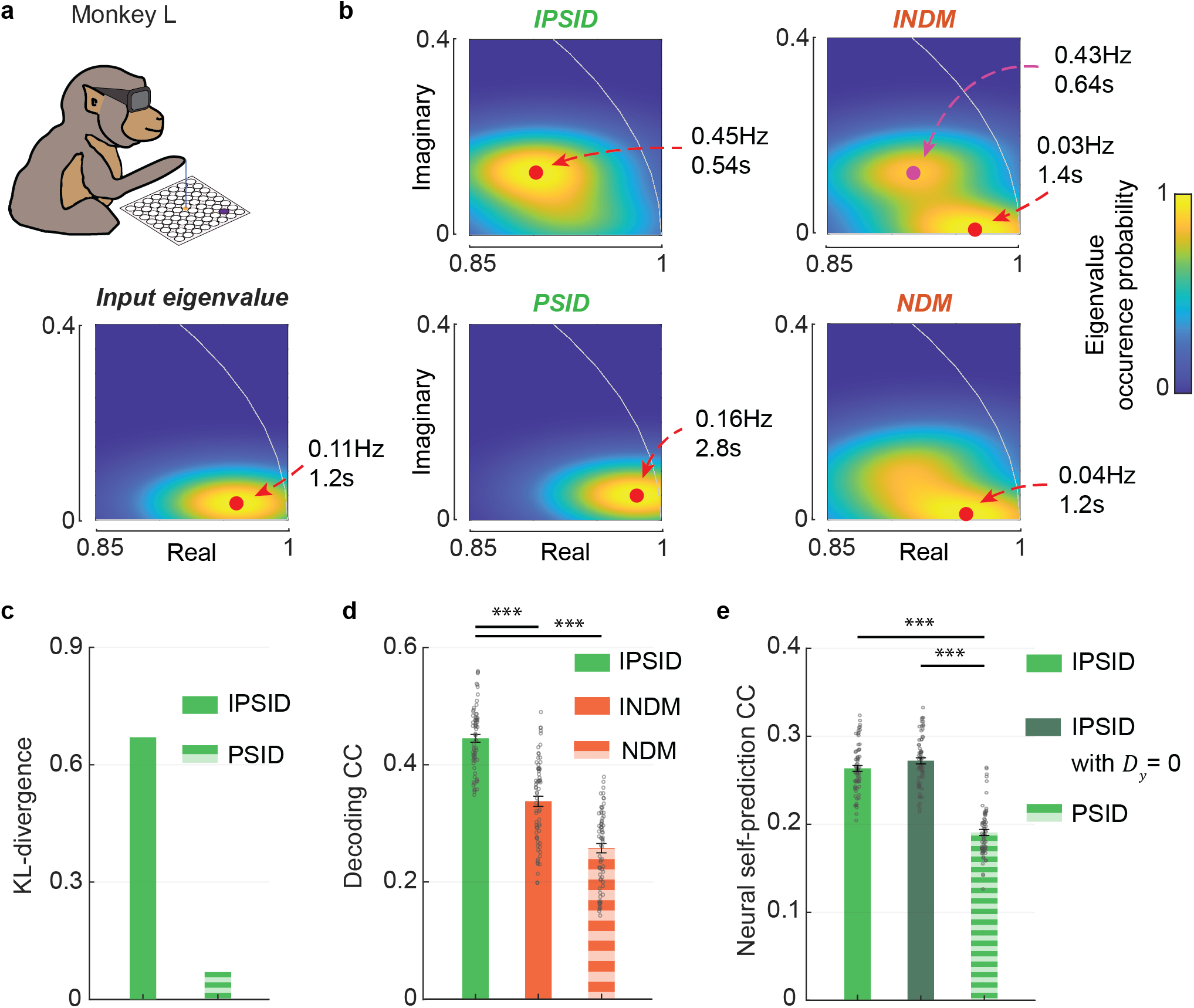
In a third monkey, IPSID again uncovers distinct and more accurate intrinsic behaviorally relevant neural dynamics in spiking activity by considering task instructions as inputs to the brain. Similar to **Fig. 6** for the third subject (monkey L, *n* = 70 cross-validation folds across 2 channel subsets and 7 recording sessions, **SI Methods**). The task is the same as in **Fig. 6** for a different first monkey while the task is different from **Fig. 7** for a different second monkey.

**Fig. S12.**
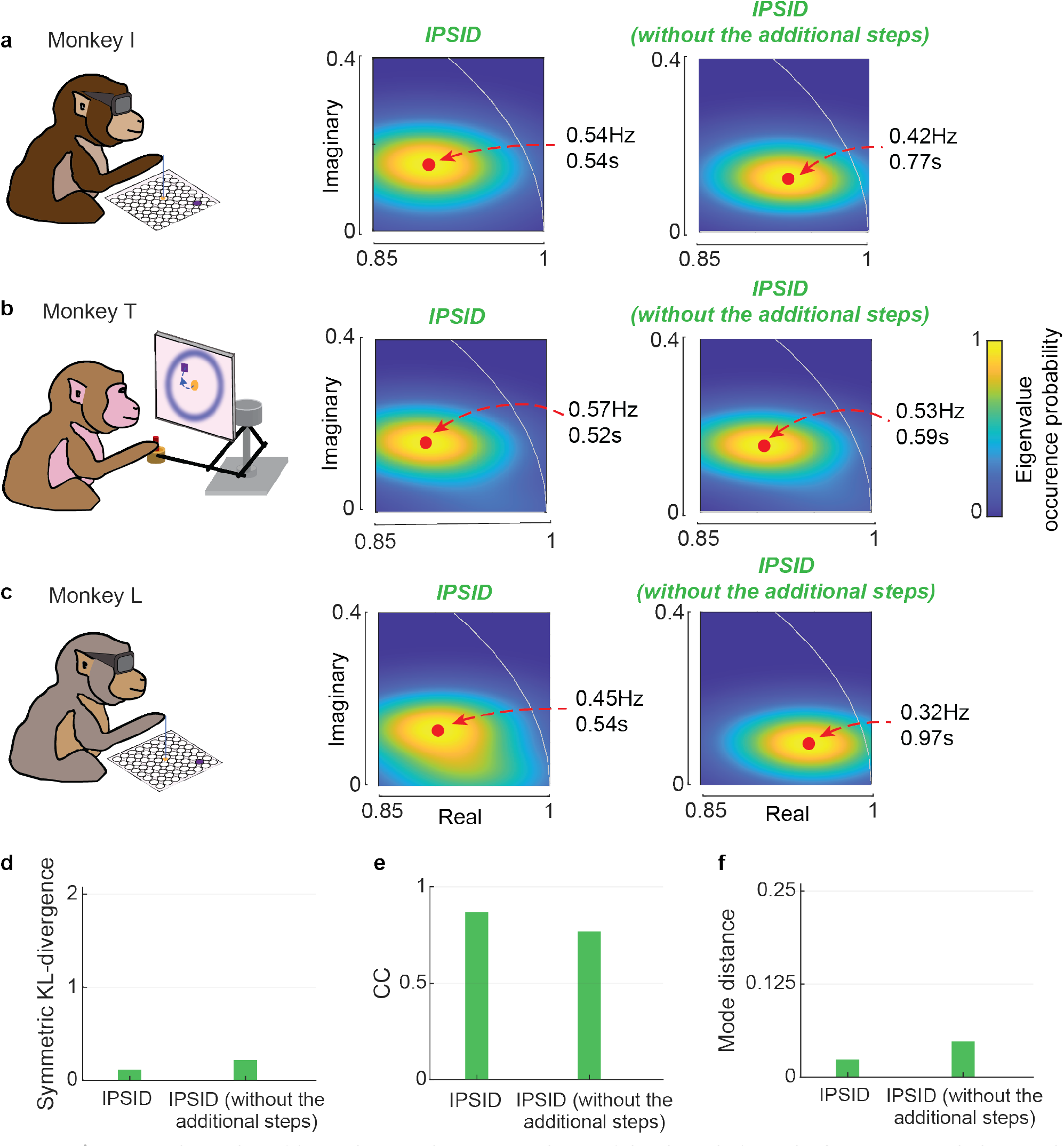
Even without the additional steps that ensure the model only include paths from input to behavior that are encoded in the recorded neural activity, IPSID finds largely similar eigenvalues across three monkeys and two tasks from two independent datasets. **(a)** Same as **Fig. 6b**, showing the eigenvalues learned for both versions of IPSID, with and without the additional optional steps (**Fig. S5** vs. **Fig. S1**, respectively). **(b-c)** Similar to (a) for the second and third subjects (**Fig. 7, Fig. S11**). Both versions of IPSID largely find similar eigenvalues. Nevertheless, to ensure only paths from input to behavior that are encoded in the recorded neural activity are included in the learned models, IPSID with the additional steps that ensures this property is used in **Figs. 6–8** and **Fig. S11**. **(d-f)** The similarity of the eigenvalues across three monkeys from two independent datasets with two distinct tasks when learned by IPSID with vs. without its additional steps as quantified by the symmetric KL-divergence, correlation coefficient, and mode distance. Notation is as in **Fig. 8d-f**. As quantified by all three metrics, both versions of IPSID find largely consistent eigenvalues across the two monkeys/tasks, with the additional steps helping IPSID reveal the similarity slightly more clearly (compare with the much larger distance/divergence and much smaller CC for INDM in **Fig. 8**).

**Fig. S13.**
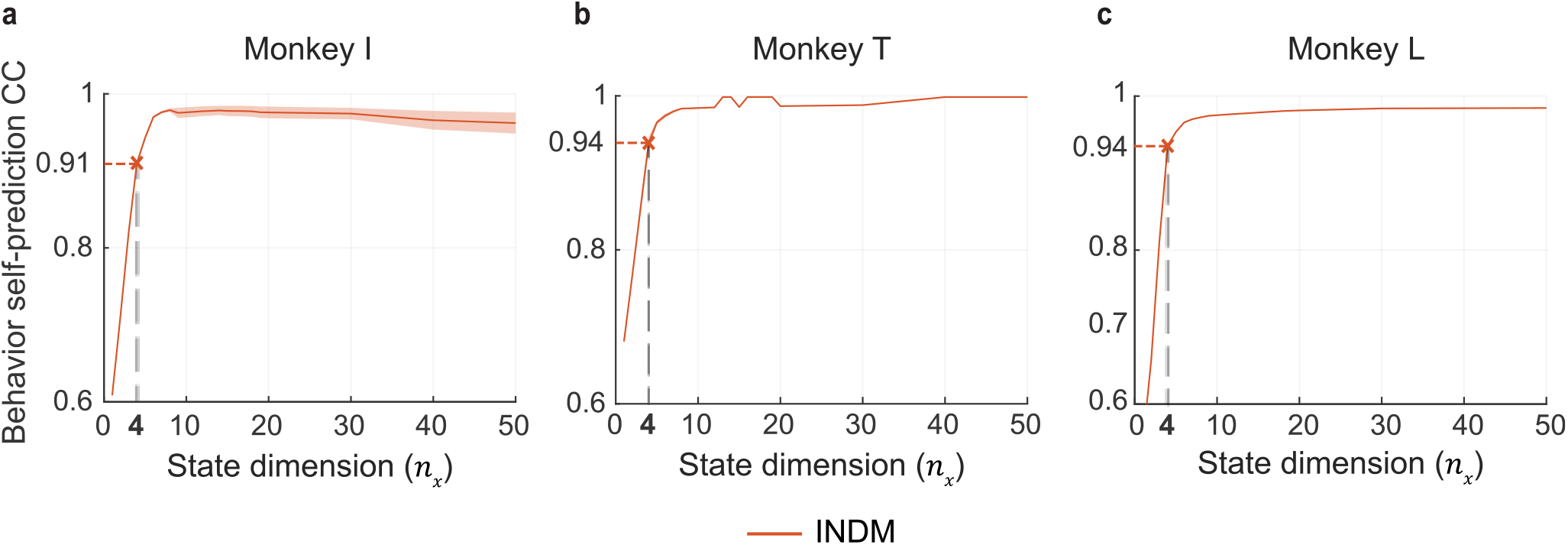
Latent state dimension of 4 is sufficient to capture most of the behavior dynamics. **(a)** Cross-validated behavior self-prediction versus latent state dimension for the first subject (monkey I) with reaches to random targets on a grid (**Fig. 6a**). Here INDM is applied to behavioral data and the learned model is used to predict the current behavior signal from its past. Using a latent state dimension of 4, behavior self-prediction CC reaches 91% of the ideal value (CC=1). **(b)** Similar to (a) for the second subject (monkey T) with sequential reaches to random targets (**Fig. 7a**). Using a latent state dimension of 4, behavior self-prediction CC reaches 94% of the ideal value (CC=1). **(c)** Similar to (a) for the third subject (monkey L) performing the same task as in (a). Using a latent state dimension of 4, behavior self-prediction CC reaches 94% of the ideal value (CC=1).

## Footnotes

† tf.keras.layers.RNN

